# ERBB2 is a Key Mediator in Hearing Restoration in Noise-Deafened Young Adult Mice

**DOI:** 10.1101/838649

**Authors:** Jingyuan Zhang, Daxiang Na, Holly J. Beaulac, Miriam Dilts, Kenneth S. Henry, Anwen Bullen, Patricia M. White

## Abstract

Noise-induced hearing loss (NIHL) affects over ten million adults in the United States, and has no biological treatment. We hypothesized that activation of signaling from ERBB2 receptors in cochlear supporting cells could mitigate cochlear damage. We adopted a new timeline for assessing mitigation that parallels hearing recovery from damage in avians. We drove expression of a constitutively active variant of ERBB2 (CA-ERBB2) in cochlear supporting cells three days after permanent noise damage in young adult mice. Between 100-200 supporting cells in the apical cochlea expressed a lineage marker, indicating competence to express CA-ERBB2. Hearing thresholds were assessed with auditory brainstem response tests, and hearing recovery was assessed over a ninety-day period. Mice harboring CA-ERBB2 capability had similar hearing thresholds to control littermates prior to noise exposure, immediately after, and 30-days after. Sixty and ninety days after noise exposure, CA-ERBB2+ mice demonstrated a partial but significant reversal of NIHL threshold shifts at one in five frequencies tested, which was in the region of CA-ERBB2 expression. We evaluated inner and outer hair cell (IHC and OHC) survival, synaptic preservation, stereociliary morphology, and IHC cytoskeletal alterations with histological techniques. Improved IHC and OHC survival were observed in the basal cochlea. No differences were seen in synaptic numbers or IHC cytoskeletal alterations, but more stereocilia may have been preserved. These data indicate, for the first time, that ERBB2 signaling in supporting cells can promote hair cell survival and partial functional recovery, and that permanent threshold shifts from noise may be partially reversed in mice.

## Introduction

Noise-induced hearing loss (NIHL) is a permanent sensory disability disproportionately affecting former military service personnel (Humes, Joellenbeck et al. 2006, Pearson 2009). Providing services and disability benefits for American veterans with NIHL costs over a billion dollars annually (Annual-Benefit-Report 2014). Noise exposure can increase auditory thresholds for hearing (NIDCD 2019), potentiate tinnitus (Eggermont and Roberts 2004), and exacerbate the effort of understanding speech in noise (Hygge, Evans et al. 2002). Cellular correlates of noise damage include the deaths of outer and inner hair cells (OHC and IHC) (Chen and Fechter 2003), broken cellular structures such as stereocilia and the tunnel of Corti (Lim and Dunn 1979), increased inflammation (Hirose and Liberman 2003), and disrupted auditory synapses (Puel, Ruel et al. 1998).

Birds readily regenerate lost auditory hair cells after noise (Corwin and Cotanche 1988, Ryals and Rubel 1988) and drug damage (Tucci and Rubel 1990). Some surviving supporting cells divide, and others trans-differentiate into *de novo* hair cells. Injured auditory hair cells and supporting cells are presumably repaired. The ability to discriminate between different sounds returns after one month (Ryals, Dent et al. 2013). A similar response has not been shown in mammals. To facilitate mammalian hair cell regeneration, multiple candidate pathways are under investigation, including NOTCH (Mizutari, Fujioka et al. 2013), FGF22 (Li, Hang et al. 2016), WNT (Shi, Cheng et al. 2010), and SHH (Lu, Chen et al. 2013).

We have focused on ERBB2 activation as a candidate signal for initiating a damage response in the cochlea. ERBB2 is a receptor tyrosine kinase in the EGFR family. In other epithelial tissues, ERBB2 is activated after stretch damage (Vermeer, Einwalter et al. 2003) to promote repair and regeneration. ERBB2 and the other three members of the EGFR receptor family are all expressed by adult mammalian supporting cells (Hume, Kirkegaard et al. 2003), as are ligands such as neuregulin (Zhang, Ding et al. 2002). Early experiments suggested that ligands for the EGFR family promoted rodent supporting cell proliferation (Yamashita and Oesterle 1995). Endogenous ERBB2 signaling in supporting cells has been shown to maintain cochlear neuronal survival through an indirect signaling cascade (Stankovic, Rio et al. 2004). To assess the effects of ERBB2 activation, we have employed genetic methods to drive expression of a constitutively active variant of ERBB2 (CA-ERBB2) in supporting cells (Zhang, Wang et al. 2018). This variant autodimerizes within the membrane to stimulate activation of PI3K (Xie, Chow et al. 1999), a key regeneration modulator in avian tissues (Montcouquiol and Corwin 2001). In previous studies of CA-ERBB2 activation in undamaged, neonatal mouse cochleae, supporting cells harboring CA-ERBB2 indirectly stimulated changes in surrounding cells, including ectopic hair cell differentiation (Zhang, Wang et al. 2018). While promising, the maturing rodent cochlea becomes less responsive to regeneration signals (Maass, Gu et al. 2015), rendering the utility of this pathway unknown.

To test the efficacy of ERBB2 signaling in hearing restoration after noise damage, we have employed the same genetic mouse model used previously (Zhang, Wang et al. 2018) in a noise damage paradigm. Young mice harboring a CA-ERBB2 transgene (Xie, Chow et al. 1999) were exposed to traumatic noise, which significantly and permanently reduces auditory function in control mice of that age and strain (Myint, White et al. 2016). The mice were subjected to pharmacological induction of the Tet-ON transgene in supporting cells (Cox, Liu et al. 2012). Functional and anatomical studies were conducted to assess the long-term effects of this intervention. Notably, we extended the period of analysis to 90 days post noise exposure (DPN), to assess improvements from permanent threshold shifts, in parallel to the time course of hearing restoration in avians.

## Materials and Methods

### Experimental Subjects

The following mouse strains were used in these experiments: *Fgfr3-iCre* (Cox, Liu et al. 2012) was a kind gift from Dr. Jian Zuo. *CA-ErbB2* mice were purchased from Jackson Laboratories. Both lines were bred to CBA/CaJ mice (Jackson Laboratories) for five generations. *ROSA-floxed-rtTA/GFP* was a kind gift from Dr. Lin Gan. This line was made using the same modifications described by others (Belteki, Haigh et al. 2005) on 129SvEv and backcrossed four times to C57BL6/J (Jackson Laboratories). When crossed to an appropriate CRE-expressing line, the young adult F1 cochlear supporting cells displayed GFP fluorescence. *Fgfr3-iCre* mice were crossed to *CA-ErbB2* mice, and then male mice harboring a single copy of both modifications were crossed to homozygous *ROSA-floxed-rtTA/GFP* females.

Both male and female mice were used equally throughout these experiments. Mice were maintained on a 12-hour light-dark cycle. No more than five adults were housed per cage with food and water available *ad libitum*. Mice also received ample nesting supplies and small houses. The day that pups were found was designated P0. Genotyping primers and protocols are available upon request.

In all, 56 mice were prepared for this study in eight separate cohorts. In the first four cohorts, 20 mice received tamoxifen (tam) injections, hearing tests, noise damage, and sequential doxycycline (dox) and furosemide (furo) injections. One mouse died immediately after the dox/furo treatment. In the fifth cohort, 60% (3 of 5) of the mice died after dox/furo injections. The surviving two mice from that cohort were excluded from the results. A sixth cohort contained 17 mice: 10 mice that received tam injections, noise exposure, and saline/furo treatments, and were aged to 120 days for no-dox controls, as well as seven mice that were processed for SEM after aging to 120 days. A seventh cohort of five mice were used to assess hair cell loss one week after noise damage. A final cohort of nine mice, also euthanized one week after noise exposure, were used for qPCR analysis.

### Ethical Approval

All experiments using animals were approved in advance by the University Committee on Animal Research (UCAR) at the University of Rochester, protocol number 2010-011, PI Patricia White.

### Experimental Design

To assess a candidate signal for hearing restoration, we took advantage of the age-dependent sensitivity to noise documented for CBA/CaJ mice (Henry 1984) and their F1 hybrids (Erway, Shiau et al. 1996, Milon, Mitra et al. 2018). Hearing thresholds were measured for each mouse by Auditory Brainstem Response (ABR) in the left ear prior to noise damage. Five frequencies were tested: 8, 12, 16, 24, and 32 kHz. We then exposed the mice to an 8-16 kHz noise band at 110 dB for 2 hours, a noise dose that we anticipated would induce large, permanent threshold shifts in young CBA/CaJ adults (Ohlemiller, Wright et al. 2000). The efficacy of the noise exposure for damaging hearing was assessed one day later. All mice had initial threshold shifts of more than 25 dB on average for all five frequencies. Hearing recovery at 30, 60, and 90 days after noise damage was also assessed by ABR. Prior to initiating experiments, it was agreed that mice with permanent threshold shifts less than 20 dB on average for all five frequencies at 30 days post noise (DPN) would not be further analyzed. We anticipated that mice with poor initial hearing (e.g. >60 dB in all frequencies) or mice that did not incur a large threshold shift from noise could present different outcomes from the others. This may be more likely due to a failure of process, such as cage movement during noise exposure. Such mice were sporadic and never siblings.

We used blinding and randomization in the following ways: The person handling the mice was blinded to genotype. At the completion of the experiment, all ABR files were masked and randomized, and scored by an independent expert (KH). Confocal images used in synaptic analysis were also masked and randomized prior to threshold setting. Mice were randomly assigned to equivalent positions in the noise exposure chamber, and post-mortem samples were randomly assigned to different kinds of analysis (cochleogram, sectioning, etc.). All randomization was performed by drawing cards from a fair deck.

### Procedures, including Materials and Equipment

#### Administration of substances to mice

Mice received three i.p. injections of tamoxifen (tam: 75 mg/kg, Sigma, #T5648) dissolved in corn oil (Sigma, #C8267) at 5 mg/kg, starting at P21. Injections were once a day, given over three consecutive days. Mice were fed chow laced with doxycycline (200 mg/kg of chow, BioServ #S3888), as described in the text. They also received a single i.p. injection of doxycycline hyclate (dox: 100 mg/kg body weight, Sigma Aldrich #D9891), which was freshly prepared as 10 mg/ml stock in 0.9% sterile saline, and two injections of 5-ethynyl-2’-deoxyuridine (EdU: 0.01 mg/kg, Invitrogen #A10044), made as a 10 mM stock solution in DMSO and diluted for injection to 40% strength in 0.9% sterile saline. The dox injection was followed by an injection of furosemide (furo: 400 mg/kg, Merck #21784-46150) dissolved in 0.9% sterile saline after a 30-minute interval. Although all the mice received EdU injections, we were unable to perform a full analysis on proliferation due to a state-mandated laboratory shutdown, and do not present this data.

#### Noise exposure

Awake P35 mice were exposed to noise, limited to the 8-16 kHz octave band at 110 decibels (dB) for 120 minutes. This noise level was anticipated to induce permanent threshold shifts in young CBA/CaJ adults (Ohlemiller, Wright et al. 2000). Mice were each placed into individual triangular wire mesh cages, 12 cm x 5 cm x 5 cm, in an asymmetric plywood box with a 2250HJ compression speaker (JBL) and 2382A biradial horn (JBL) mounted on top. This apparatus was contained within a sound booth. The speaker was driven by a TDT RX6 multifunction processor and dedicated attenuator, and controlled with TDT RPvdsEx sound processing software. The sound level was calibrated with a Quest Technologies sound meter, Model 1900. Mice were exposed between the hours of 9 AM and 3 PM to control for circadian rhythm effects (Meltser, Cederroth et al. 2014), and the sound level was checked with an iPhone using the Faber Acoustical SoundMeter app and the iPhone’s internal microphone each morning before use. The iPhone was previously calibrated with the SoundMeter app software and the solid state 94 dB source (provided with the Quest sound meter) in a sound booth.

#### Auditory testing

Mice were tested on the schedule described in the text. Closed field auditory testing was conducted using a Smart EP Universal Smart Box (Intelligent Hearing Systems) with a ED1 speaker (Tucker Davis Technologies). The apparatus was calibrated with a Qwest sound meter less than one month prior to the start of the experiment. Mice were anesthetized with an intraperitoneal injection of ketamine (80 mg/kg) in a sterile acepromazine/saline mixture (3 mg/kg). Speaker outputs were bundled with a 10B+ OAE microphone, and placed at the opening to the external auditory meatus.

Auditory brainstem response (ABR) stimuli were 5-ms clicks, or 5-ms tone pips presented at five frequencies between 8 and 32 kHz. Stimuli began at 75 dB amplitude and decreased by 5 dB steps to 15-25 dB. 512 sweeps were averaged for each frequency and amplitude. Electrical responses were measured with three subdermal needle electrodes (Grass): one inserted beneath each pinna, and a third placed at the vertex. ABR thresholds for a particular frequency were determined by any part of the last visible trace (dB). If no waveform was observed, the threshold was recorded as “80 dB.” As we did not measure responses to stimuli greater than 75 dB, the hearing loss from noise may have been underestimated. Wave I amplitude and latency measurements were made using IHS software capabilities.

#### Tissue preparation for immunostaining

Cochlear organs were dissected out of freshly euthanized animals. The stapes were removed and a hole was made in their apical tips to allow for adequate fluid exchange. Tissues were immersed in 4% paraformaldehyde in PBS for at least overnight. Tissues were decalcified in 0.1M EDTA at 4°C on a rotating platform for four days. For cryosectioning, tissues were immersed in 30% sucrose in PBS overnight, embedded in OCT, frozen in liquid nitrogen, and cryosectioned at 20 microns. For hair cell (HC) counts and synapse analysis, whole mount preparations were microdissected into turns as previously described (Montgomery and Cox 2016), mapped using the ImageJ plug-in from Massachusetts Eye and Ear Infirmary, and immunostained.

#### Antibodies

The following primary antibodies were used: mouse anti-CTBP2 (aka C-Terminal Binding Protein 2; 1:200; BD Transduction Laboratories; RRID:AB_399431), chicken anti-GFP (1:500; Abcam; RRID:AB_300798), rabbit anti-GLAST (aka EAAT1; 1:100; Abcam; RRID:AB_304334), mouse anti-GRIA2 (aka GluR2/GluA2; 1:2000; Millipore; RRID:AB_2113875), mouse anti-MAP2 (aka Microtubule-Associated Protein 2; 1:250; R&D Systems, Cat #: MAB8304, RRID:AB_2814693), rabbit anti-Myosin7 (MYO7) (1:200; Proteus; RRID:AB_10013626), mouse anti-Myosin7 (1:100; Developmental Studies Hybridoma Bank, RRID:AB_2282417), and goat anti-Oncomodulin (OCM) antibody (1:1000; Santa Cruz; RRID:AB_2267583). The following secondary antibodies, all purchased from Jackson ImmunoResearch, were used: Donkey anti-Chicken 488 (1:500; RRID:AB_2340376), Donkey Anti-Mouse 488 (1:500; RRID:AB_2340849), Donkey Anti-Rabbit 594 (1:500; RRID:AB_2340622), Donkey Anti-Rabbit 647 (1:200; RRID:AB_2340625), Donkey Anti-Goat 647 (1:200; RRID:AB_2340438), Alexa 594 Goat Anti-Mouse (IgG1, 1:500; RRID:AB_2338885), Alexa 488 Goat Anti-Mouse (IgG2a, 1:500; RRID:AB_2338855), Alexa 594 Goat Anti-Mouse (IgG3, 1:500; RRID: AB_2632543), Alexa 488 Goat Anti-Mouse (IgG1, 1:500; RRID: AB_2632534).

#### Immunostaining

For immunostaining on mouse sections, blocking was carried out at room temperature for two hours in 1% Tween / 5% goat serum (Jackson) in Tris-buffered saline. Antibody incubations were carried out overnight at 4°C in block solution. For whole mount staining, dissected mapped turns were flash frozen in liquid nitrogen, allowed to thaw, and washed in room temperature Dulbecco’s PBS (Gibco). Prior to immunostaining, tissues were blocked for one hour in 1% Triton / 5% donkey serum in PBS. Primary antibody incubations of anti-MYO7, anti-OCM, anti-GFP, anti-CTBP2, and anti-GRIA2 were performed at 37°C for 20 hours. The tissue was washed in PBS, and secondary antibody incubation was performed at 37°C for an additional 2 hours. All tissue was mounted using ProLong Gold (Fisher). Whole mounts were placed between two 50 mm coverslips for imaging.

#### Confocal microscopy and image processing for figures

Imaging was performed on an Olympus FV1000 or a Nikon A1R HD, both laser-scanning confocal microscopes. ImageJ (NIH) was used to Z-project maximal brightness in confocal stacks, select optical sections, and combine colors. Photoshop CS5 (Adobe) was used to set maximal and background levels of projections for the construction of figures. Composite images for cochleograms were assembled in Photoshop by pasting each optical section into its own layer, and merging the pieces of the optical sections where HCs were evident. Alternatively, projections of confocal stacks were used when individual IHCs could be clearly distinguished. Composite or projected images for the mapped regions of each organ were assembled in a single file in Photoshop. 100-micron lengths were pasted onto the images on the row of pillar cells, and OHCs were counted to determine the percent lost. Any lengths containing post-mortem damage, e.g. no sensory epithelium, were omitted from the analysis. In the scatter plot, such regions are left blank. Where the loss of OHCs was great enough (>30%), an average OHC count, determined from low-frequency regions of the same cochlea, was used as the denominator (Viberg and Canlon 2004).

#### 3-D reconstructions of synaptic components

To quantitatively assess synaptic number and distribution in IHCs, we used immunofluorescence and confocal microscopy in conjunction with three-dimensional modeling. Anti-CTBP2 labeled IHC nuclei and pre-synaptic ribbon structures (Schmitz, Konigstorfer et al. 2000), and anti-GRIA2 was specific for the post-synaptic AMPA receptor (Usami, Matsubara et al. 1995). Anti-MYO7 labeled IHCs (Hasson, Heintzelman et al. 1995). The most effective imaging of ribbon synaptic complexes had illumination bathing the neurites, with light collection from the stereociliary side. Confocal data files (.oib) of appropriately stained IHCs, imaged at 200X on the FV1000, were obtained for the entire dataset. These files were blinded and randomized using a deck of cards. We imported the data into Amira 6.0 (Visualization Science Group) to generate 3-D reconstructions.

A Myo7a isosurface was created, appropriately thresholded, and exported. The CTBP2 and GRIA2 datasets were subjected to blind deconvolution, performed using the “maximum likelihood estimation” package for Amira. Briefly, point spread functions used for deconvolution were calculated using the optical parameters of the microscope (N.A. =1.40, UPlanSAPO 100x oil objective, 400-800 nm) and the wavelength of interest; the datasets were deconvolved for 5 iterations. The data were despeckled, followed by isosurface creation with appropriate thresholding. The Connected Components function was used to output a complete list of all CTBP2 structures. To identify pairs of synaptic components, we used the Amira XImagePAQ AND functionality, which identifies adjacent staining elements. Those elements were dilated and subtracted from the CTBP2 dataset to identify “orphan” or unpaired synaptic ribbons. The complete list and the “orphan” list were compared to obtain paired ribbons. Only after synaptic pairs were processed was the MYO7 data visualized for display.

#### Quantitative PCR

Mice were exposed to noise, injected with doxycycline and furosemide three days later, and euthanized seven days after noise exposure. Cochleae were isolated and total RNA was extracted using the RNeasy mini kit (Qiagen). cDNA was generated using the qScript cDNA Synthesis kit (QuantaBio). Total RNA concentration and purity was determined with the NanoDrop 1000 spectrophotometer (NanoDrop) and RNA quality was assessed with the Agilent Bioanalyzer (Agilent). cDNA was prepared and amplified from RNA samples with the Ovation PicoSL WTA System v2 (NuGEN Technologies), following manufacturer’s recommendations. Quantitative RT-PCR was performed on all samples using 12ng of amplified cDNA per reaction with TaqMan assays and TaqMan® Universal PCR Master Mix, No AmpErase® UNG on the QuantStudio 12K Flex™ Real-Time PCR System (Applied Biosystems). Comparative ΔΔCt was performed to determine relative mRNA expression, where relative abundance of each gene was first internally normalized to the geometric mean Ct for the reference genes, Gapdh and Hprt1. Quantitative RT-PCR and primary data analysis was performed by the Genomics Research Center at University of Rochester. TaqMan assays used for this analysis were: rat Erbb2 (Rn01461176_g1), mouse Erbb2 (Mm01306784_g1), Erbb3 (Mm01159999_m1), Erbb4 (Mm01256793_m1), Egfr (Mm01187858_m1), Atoh1 (Mm00476035_s1), Nt3 (Mm01182924_m1), Hprt (Mm03024075_m1), Gapdh (Mm99999915_g1).

#### Scanning electron microscopy

Cochleae were rapidly dissected out of the cranial bone one animal at a time to minimize the amount of time between death and fixation at room temperature. 400 µl of fixative, containing 4% paraformaldehyde and 2% gluteraldehyde in 0.1M sodium cacodylate buffer, was gently pipetted through the open oval window, exiting through the hole made in the apical tip. Tissues were post-fixed at 4°C on a rotating platform overnight, rinsed three times with distilled water, decalcified in 10% EDTA in 100 mM Tris pH 7.4 for three days, and then rinsed again. Cochlea turns were dissected, post-fixed in 1% osmium tetroxide for two hours at room temperature, and then processed through the thiocarbohydrazide-osmium tetroxide repeated procedure (Davies and Forge 1987). Samples were then dehydrated, critical point dried, mounted on support stubs with silver paint, and sputter coated with platinum. Imaging by secondary electron detection was carried out on a JEOL JSM6700F scanning electron microscope, operating at 1.2 and 5kV.

### Statistical Analysis and Data Reporting

Statistical analyses were performed in R (R Core Team). Datasets that failed the Shapiro-Wilk normality test included some of the wave I amplitude and latency measurements, hair cell losses for the cochleograms, where noise-damaged CA-ERBB2 cochleae were compared to noise-damaged control cochleae, and synaptic sizes at 12 and 24 kHz, where noise-damaged CA-ERBB2 cochleae were compared to both noise-damaged control cochleae and no-noise control cochleae. Non-normal amplitude and latency groups were analyzed with Wilcoxon rank sum test with the continuity correction. Cochleogram data were analyzed with the Kruskal-Wallis rank sum test, and pairwise comparisons of synaptic sizes were adjusted for multiple comparisons with the Bonferroni method. ABR thresholds for noise-damaged CA-ERBB2 mice and noise-damaged control mice were compared with a multivariable ANOVA. All subsequent pairwise analysis was performed using Student’s two-tailed t-test with the Bonferroni adjustment for multiple comparisons. Wave I latency and amplitude sets that passed the Shapiro-Wilk normality test were analyzed with the Welsh two sample t-test. Statistical significance was deemed for p-values less than 0.05. Exact p values are reported, except for those with values less than 2.2 × 10^-16^, which is the lowest value R can calculate.

A post-hoc power analysis was conducted on the variance of threshold recovery (ABR) for control mice. For all five frequencies, the average recovery was 1.6 dB and the pooled variance was 11.24 dB. For a sample size of nine experimental mice, we have 80% confidence in results that differ by more than 10.5 dB.

All primary data were reviewed by the senior author prior to the assembly of figures for this paper. Any differences between the data presented here and any other public presentation are due to errors that were caught in that review. All the original data generated for this article will be made available prior to publication through UR Research.

## Results

### Principle of the noise/drug hearing damage mouse model to study the role of ERBB2 in hearing recovery

We hypothesized that activation of ERBB2 signaling in cochlear supporting cells after noise damage would promote hearing recovery. To test this hypothesis, we generated mice harboring a Tet-ON constitutively active rat ERBB2 transgene (Xie, Chow et al. 1999), a floxed ROSA-rtTA-GFP knock-in, and the inducible Fgfr3-iCRE gene (Fig. 1A, pink box). We refer to these mice as “CA-ERBB2” mice, and their littermates harboring two of the three modifications as “control” mice (Fig. 1A, gray box). Activation of iCRE at P21-P23 in supporting cells was assessed with the GFP lineage marker (Fig. 1B, B’, green). We counted 28 ± 8 GFP+ cells/mm in the apical third of the cochlea, where 93% of the GFP+ cells resided. 64% of the GFP+ cells were located under the OHC layer with a morphology consistent with a Deiters’ cell identity (Fig. 1B’, inset, cf. red and green. A cartoon is provided below the side view). The remaining cells were in positions consistent with pillar cells (8%) or Hensen’s cells (27%). No GFP immunoreactivity was observed in mice harboring the floxed ROSA rtTA gene and the ERBB2 transgene, but lacking Fgfr3-iCRE (Fig. 1C, green).

**Figure 1.**
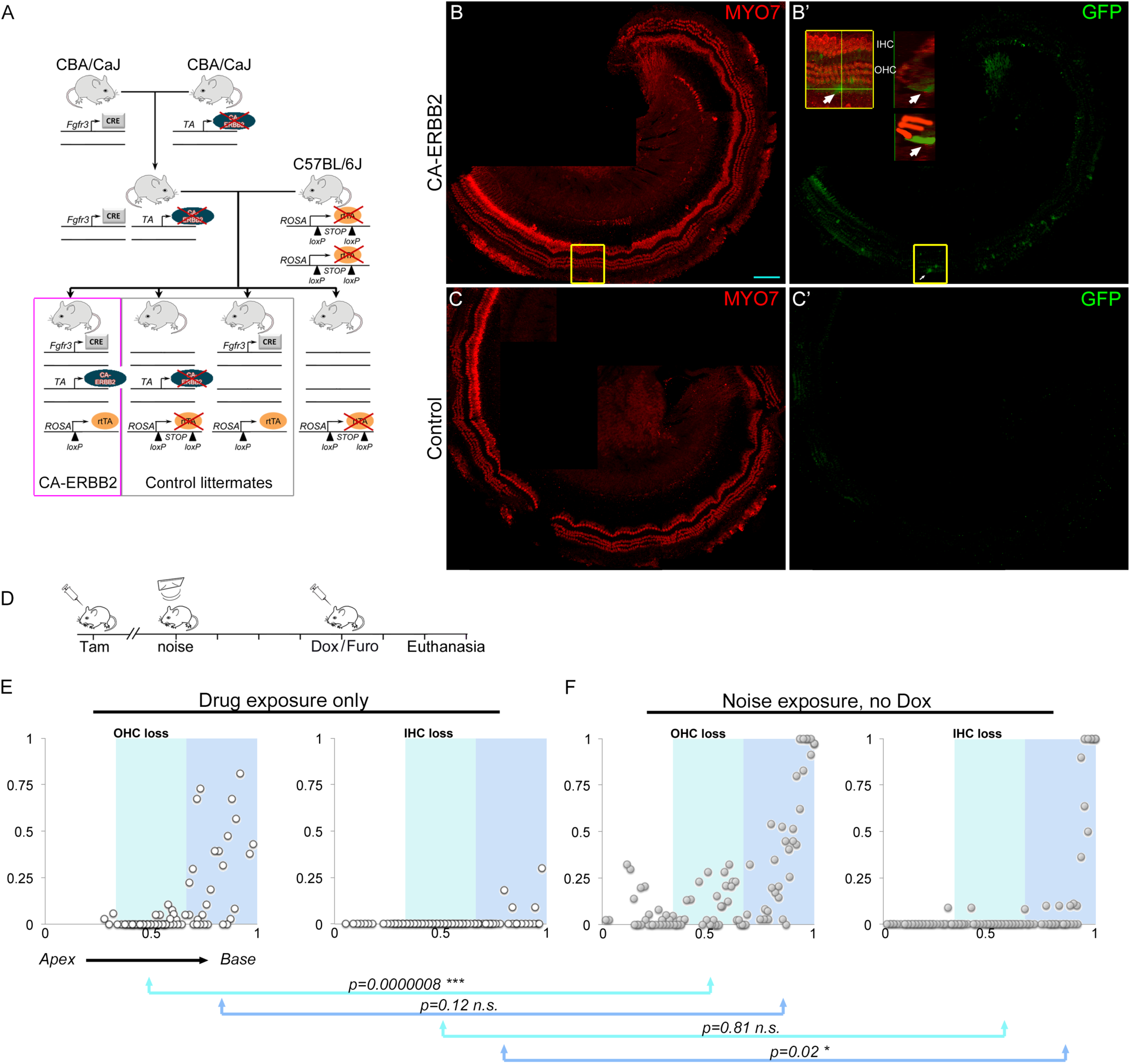
Experimental design, GFP+ cell distribution, and modes of hair cell damage. A. Breeding strategy diagram. Genes that are not expressed in a particular genetic combination are covered with a red “x”. Mice harboring the *Fgfr3-iCre* and *CA-ERBB2* modifications were maintained separately on a CBA/CaJ background. Mice heterozygous for both genes were crossed with C57BL/6J mice homozygous for *ROSA-floxed-rtTA-IRES-GFP*. For the progeny of the second cross, the four possible genotype combinations are shown. Mice harboring both *ROSA-floxed-rtTA-IRES-GFP* and *Fgfr3-iCRE* recombine the *loxP* sites in Fgfr3-iCRE expressing cells after the injection of tamoxifen, enabling the production of TA protein and GFP. Upon doxycycline treatment, TA protein can drive expression of CA-ERBB2. Mice harboring all three modifications are called “CA-ERBB2 mice” (pink box), whereas mice harboring two out of three modifications are called “control mice” (gray box). B, B’. MYO7a (red) and GFP (green) immunoreactivity at P35 in the apical turn from a CA-ERBB2 mouse after tamoxifen injection, showing scattered GFP+ supporting cells. The yellow box shows the location of the inset (B’), where the side panel shows a GFP+ Deiters’ cell (inset, white arrow). The cartoon below the side view highlights the morphology of three OHCs and the GFP+ cell. This mouse had 20 cells/mm in the apical turn, with only occasional cells elsewhere. GFP+ cells are competent to express CA-ERBB2. Size bar: 100 microns. C, C’. Matched image of a control cochlea without *Fgfr3-iCRE* stained for MYO7a (red) and GFP (green), to demonstrate specificity. D. Timeline of drug treatments and noise exposures for cochleograms shown in (E, F). Tam: tamoxifen; pre-test: ABR hearing test; noise: noise treatment (F); 1 DPN: ABR hearing test; Dox: doxycycline; EdU: ethynyl deoxyuridine; Furo: furosemide. Euthanasia was performed 48-72 hours after Dox/Furo injections. E. OHC loss (left panel) and IHC loss (right panel) for mice treated with tamoxifen, doxycycline and furosemide. The cochlea is segmented into the apical (4-16 kHz region, white), middle (16-32 kHz region, cyan) and basal (32-64 kHz region, blue) thirds of the cochlea. Hair cell losses were quantified in 100-micron segments along the length of each mapped organ (n=3 cochleae), revealing basal OHC loss from the drug treatments (blue, left panel). F. Same representation of hair cell loss in mice injected with tamoxifen, exposed to noise, and then injected with saline followed by furosemide (n=2 cochleae). This strategy reveals hair cell loss due to noise exposure without confounding losses due to doxycycline. Significantly more middle OHC loss is observed (p = 8 × 10^-7^, cyan, n=2-3 cochleae, Kruskal-Wallis rank sum test with Bonferroni adjustment). Significant IHC loss from noise is observed in the basal cochlea (p=0.02, blue, n=2-3 cochleae, Kruskal-Wallis rank sum test with Bonferroni adjustment).

We were unable to directly observe phosphorylation of ERBB2 by immunofluorescence in adult cochlear supporting cells, as in previous studies (Zhang, Wang et al. 2018). However, qPCR analysis for the rat ERBB2 transgene showed that level of this transcript was 8.9 ± 2.9 fold higher in CA-ERBB2 cochleae compared to control cochleae, four days after dox and furosemide administration (S. Fig. 1, p=0.027, n=4-5 cochleae, student’s two-tailed t-test with Bonferroni correction). We tested for changes in two potential downstream targets, *Atoh1* and *Ntl3*, as well as for changes in *Egfr*, mouse *Erbb2*, *Erbb3*, and *Erbb4* transcripts. No differences were seen in the levels of those genes (S. Fig. 1).

Both noise (Hamernik, Henderson et al. 1974) and dox/furo injections (Walters and Zuo 2015) have been reported to cause OHC losses. We used the schedule shown in Fig. 1D to illustrate the effects of these two treatments. Two CA-ERBB2 mice were exposed to noise at P35, and three control littermates were placed in the apparatus for the same length of time with the sound turned off. At P39, the three unexposed control mice received dox/furo injections, and the two CA-ERBB2 mice exposed to noise received saline/furo injections. All mice were euthanized 2-3 days after the injections, when hair cell loss from dox has been reported to be complete. Their cochleae were assessed for OHC and IHC survival with immunostaining for OCM and MYO7 respectively (see Methods). Mice injected with dox/furo exhibited 27 ± 13% reduction in OHCs in the basal third only of the cochlea (Fig. 1E, left panel) and little or no IHC loss (Fig. 1E, right panel), which is comparable but less severe than the previous report. CA-ERBB2 mice exposed to noise and furo, but not treated with dox, had more OHC loss in the middle third of the cochlea (cf. Fig. 1E left panel, cyan, to Fig. 1F left panel, cyan, p=8 x 10^-7^, 2-3 cochleae per condition, Kruskal-Wallis rank sum test adjusted for multiple comparisons) but not in the basal third (cf. Fig. 1E left panel, blue, to Fig. 1F left panel, blue, p=0.12, 2-3 cochleae per condition, Kruskal-Wallis rank sum test adjusted for multiple comparisons). Noise alone induced more IHC loss only in the basal third of the cochlea (cf. Fig. 1E right panel, blue, to Fig. 1F right panel, blue, p=0.02, 2-3 cochleae per condition, Kruskal-Wallis rank sum test adjusted for multiple comparisons). These data are from a small number of mice, but illustrate the kinds of hair cell losses anticipated from the damage model. We will refer to it as a mixed drug/noise paradigm.

To verify functional damage from the treatment paradigm, we recorded Auditory Brainstem Responses (ABR) to tone pips of different frequencies (Fig. 2A, “pre-test”) in cohorts of P28 CA-ERBB2 mice and control littermates. No differences were seen between the genotypes at all 5 frequencies (Fig. 2B, 8, 12, 16, 24 and 32 kHz, p=0.7, 10 control and 9 CA-ERBB2 mice, two-way ANOVA). These mice received treatments according to the schedule shown in Fig. 2A. Approximately one week after the pre-test, all mice were exposed to traumatic noise, as described above. Significant threshold shifts were observed 1-day post noise exposure (DPN, cf. Fig. 2B and 2C, p<2.2 x10^-16^, two-way ANOVA for each genotype). No differences were observed between CA-ERBB2 mice and their control littermates (Fig. 2C, p=0.91, 10 control and 9 CA-ERBB2 mice, two-way ANOVA). Two days after this hearing test, mice were fed chow laced with doxycycline (Fig. 2A, pink bar). They also received dox/furo injections (Fig. 2A). Effects of this intervention on ABR thresholds were assessed at 30, 60 and 90 DPN (Fig. 2A), by an independent expert blinded to both genotype and time point. Since threshold shifts are evaluated from before doxycycline administration, any hearing recovery constitutes restoration after NIHL.

**Figure 2.**
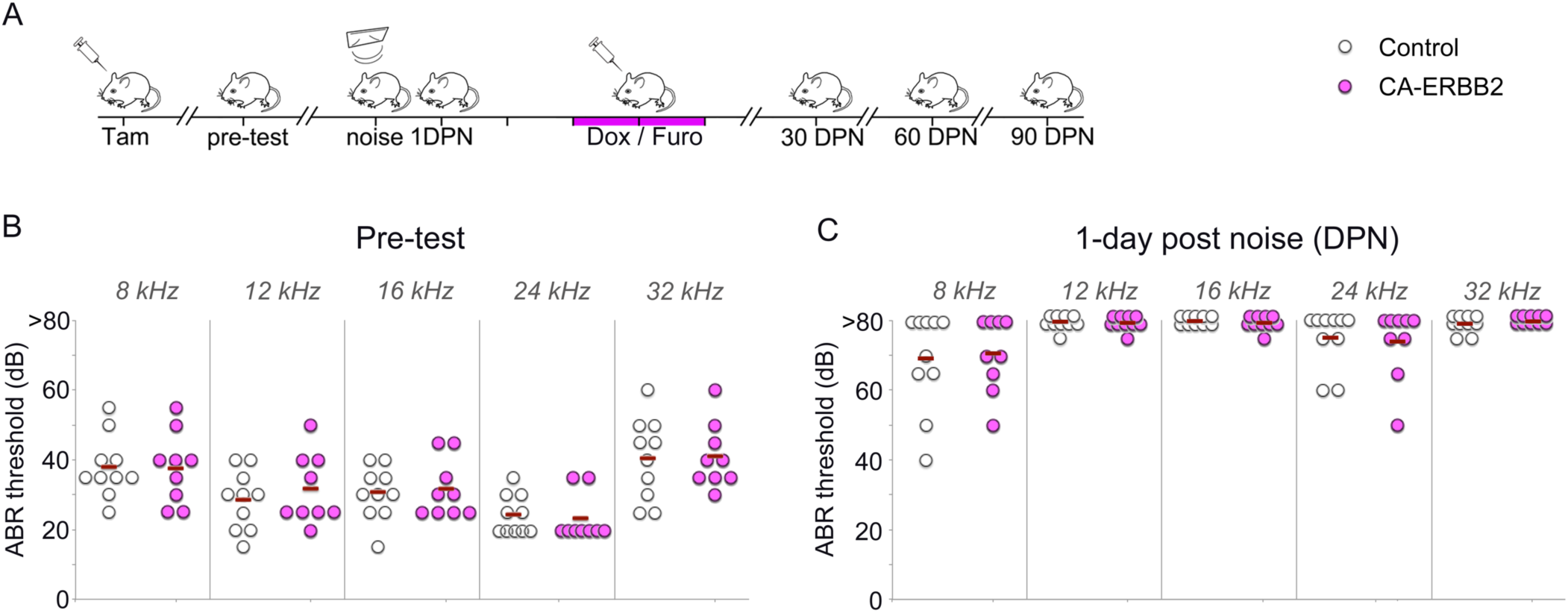
ABR threshold shifts after noise damage. A. Schematic of experimental manipulations. Injections are indicated by syringes and noise exposure by a speaker. ABR hearing tests were performed prior to noise exposure (pre-test), 1-day post noise (DPN), 30 DPN, 60 DPN and 90 DPN. Tam: tamoxifen; Dox: doxycycline; Furo: furosemide. The two-day span marked with a pink bar represents dox feeding in addition to the injection. B. ABR thresholds (in dB) are plotted for 10 control (open circle) and 9 CA-ERBB2 (pink circle) mice prior to noise exposure. Means are indicated by red bars. Five frequencies were tested. Two-way ANOVA indicates no differences between genotypes (p=0.7). C. ABR thresholds for the same mice at the same frequencies 1 DPN. If no waveform was detected, the threshold was marked as “80 dB.” Two-way ANOVA indicates no differences between genotypes (p=0.91).

### CA-ERBB2 induces partial hearing recovery in transgenic mice

To assess hearing recovery, we calculated ABR threshold shifts for each mouse by subtracting pre-test auditory thresholds from those obtained at different times after noise exposure. Positive values indicate hearing loss. In Fig. 3, ABR shifts are plotted for the 8 kHz frequency for 10 control mice (Fig. 3A, open circles) and 9 CA-ERBB2 mice (Fig. 3A, pink circles) for the four time points after noise exposure. No differences were seen overall between the genotypes at 30 DPN (p=0.33, 10 controls, 9 CA-ERBB2 mice, two-way ANOVA) or in 8 kHz ABR shifts specifically (p=0.26, 10 controls, 9 CA-ERBB2 mice, two-tailed t-test with Bonferroni adjustment). In contrast, at 60 DPN, CA-ERBB2 mice displayed partial but significant reductions in their ABR shifts (p=0.0015, 10 controls, 9 CA-ERBB2 mice, two-way ANOVA), which was due to changes in CA-ERBB2 shifts at 8 kHz (Fig. 3A, p=0.008, 10 controls, 9 CA-ERBB2 mice, two-tailed t-test with Bonferroni adjustment). This functional recovery from noise damage was sustained through 90 DPN (Fig. 3A, p=0.017, 10 controls, 9 CA-ERBB2 mice, two-tailed t-test with Bonferroni adjustment). In absolute terms, the mice recovered an average of 13.3 ± 4.6 dB, or about 41% of the threshold shift.

**Figure 3.**
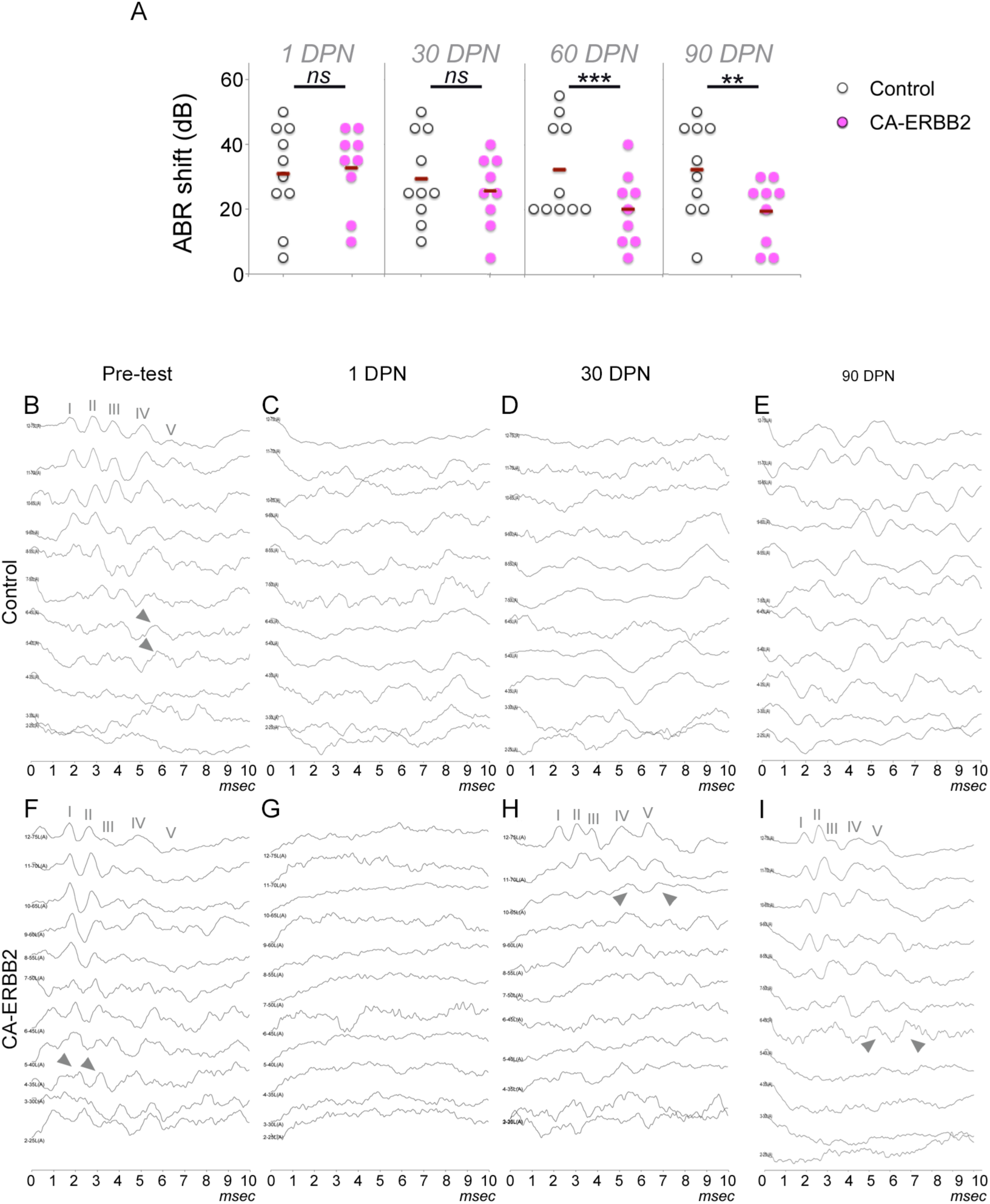
Partial, delayed recovery observed in CA-ERBB2 mice at 8 kHz. A. ABR shifts (dB) were calculated for each mouse by subtracting original ABR thresholds from those obtained at different times after noise exposure. Positive values indicate hearing loss. ABR shifts from 10 control mice (open circles) and 9 CA-ERBB2 mice (pink circles) are plotted along with means for each group (red bars). ABR shifts at 30 days were not significantly different between genotypes (two-way ANOVA p=0.34). Significant recovery was seen in the CA-ERBB2 group at 60 and 90 days (two-way ANOVA p=0.0015, 0.017, respectively), but only for 8 kHz (Bonferroni-adjusted two tailed t-tests p=0.008, 0.013, respectively). B-E. 8 kHz ABR traces for a representative control mouse are arranged in descending amplitudes of 5 dB steps, starting at 75 dB and ending at 25 dB. The x-axis indicates time in milliseconds. Peaks I-V are noted, and gray arrowheads indicate the last part of the trace that defines the threshold. ABR traces include tests performed before noise damage (B, Pre-test), 1 DPN (C), 30 DPN (D), and 90 DPN (E). No ABR waveforms are observed any time after noise damage. F-I. 8 kHz ABR traces from a representative CA-ERBB2 mouse with partial recovery at the same time points. Peaks I-V are indicated, as in (B), and gray arrowheads indicate the last part of the trace that defines the threshold. Significant improvement is observed between 30 and 90 DPN.

Representative 8 kHz ABR traces for a control mouse (Fig. 3B-E) and a CA-ERBB2 mouse (Fig. 3F-I) illustrate this hearing recovery. These mice displayed similar pre-test thresholds (Fig. 3B, cf. F, gray arrowheads) and a complete loss of hearing thresholds 1 DPN (Fig. 3C, cf. G). Hearing recovery was evident in the CA-ERBB2 mouse at 30 DPN (Fig. 3H, gray arrowheads) which continued to improve over the next two months (Fig. 3I, gray arrowheads). No such recovery was evident in the control mouse (Fig. 3E).

We also examined wave I amplitudes and latencies at 8 kHz for CA-ERBB2 and control mice before and after noise exposures (Fig. 4). Here we found that prior to noise exposure, CA-ERBB2 littermates had slight but significant increases in wave I amplitudes (Fig. 4A, p=0.008, 10 controls, 9 CA-ERBB2 mice, Wilcoxon rank sum test with continuity correction), but no differences in wave I latency (Fig. 4B, p=0.46, 10 controls, 9 CA-ERBB2 mice, Wilcoxon rank sum test with continuity correction). No differences in wave I amplitude were seen between genotypes at 30 DPN (Fig. 4C, p=0.71, 10 controls, 9 CA-ERBB2 mice, Wilcoxon rank sum test with continuity correction) or at 90 DPN (Fig. 4E, p=0.24, 10 controls, 9 CA-ERBB2 mice, Welch Two Sample t-test). However, wave I latencies for CA-ERBB2 mice were slightly but significantly delayed at 30 DPN (Fig. 4D, p=0.049, 10 controls, 9 CA-ERBB2 mice, Wilcoxon rank sum test with continuity correction) and at 90 DPN (Fig. 4F, p=0.021, 10 controls, 9 CA-ERBB2 mice, Welch Two Sample t-test).

**Figure 4.**
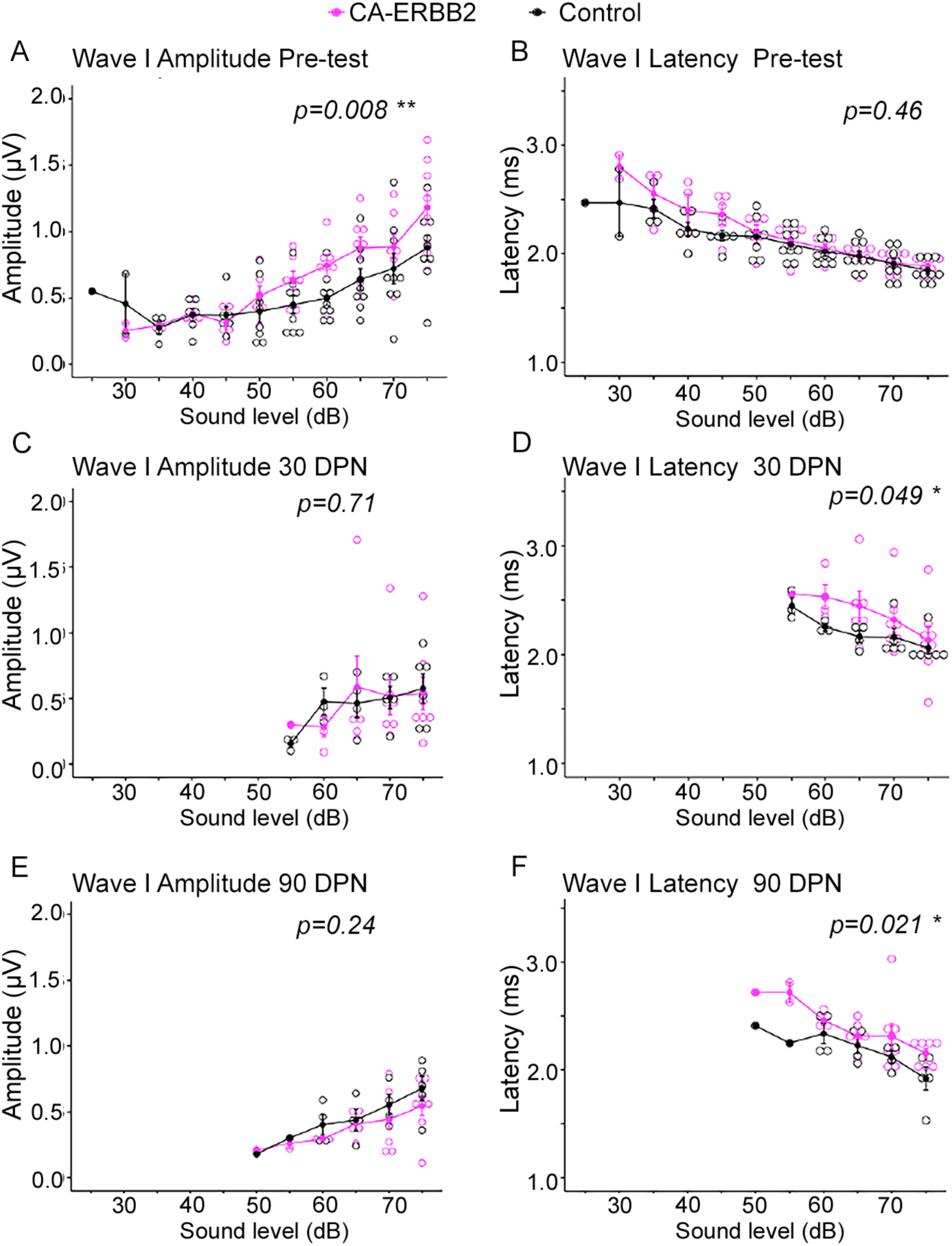
Wave I is slightly but significantly delayed in CA-ERBB2 mice at 8 kHz after hearing restoration. A. Wave I amplitudes at 8 kHz for both CA-ERBB2 (pink) and control littermates (black) before noise exposure. CA-ERBB2 mice had slightly but significantly higher amplitudes compared to control mice (p=0.008, n=10 controls and 9 CA-ERBB2, Wilcoxon rank sum test with continuity correction). B. Wave I latencies at 8 kHz are not different between CA-ERBB2 (pink) and control littermates (black) before noise exposure (p=0.46, n=10 controls and 9 CA-ERBB2, Wilcoxon rank sum test with continuity correction). C. At 30 DPN, wave I amplitudes at 8 kHz do not differ between CA-ERBB2 (pink) and control littermates (black, p=0.71, n=10 controls and 9 CA-ERBB2, Wilcoxon rank sum test with continuity correction). D. At 30 DPN, CA-ERBB2 mice (pink) have slightly but significantly longer Wave I latencies at 8 kHz compared to control littermates (black, p=0.049, n=10 controls and 9 CA-ERBB2, Wilcoxon rank sum test with continuity correction). E. At 90 DPN, wave amplitudes at 8 kHz are not different between CA-ERBB2 (pink) and control littermates (black, p=0.24, n=10 controls and 9 CA-ERBB2, Wilcoxon rank sum test with continuity correction). F. At 90 DPN, CA-ERBB2 mice (pink) have slightly but significantly longer Wave I latencies at 8 kHz compared to control littermates (black, p=0.021, n=10 controls and 9 CA-ERBB2, Wilcoxon rank sum test with continuity correction).

8 kHz was the only frequency that showed changes across time. Threshold shifts for the other four frequencies were not significantly different between genotypes (Fig. 5). Changes in threshold shift at 8 kHz was dependent on dox administration, as CA-ERBB2 mice that received saline injections instead of dox injections showed no improvement (Fig. 5E). This experiment was performed with a later cohort of mice, which had higher average thresholds at 8 kHz compared to the previous cohorts (58 dB for no dox controls vs. 38 dB for the first four cohorts). A ∼13 dB improvement to 58 dB would still be measurable for these mice.

**Figure 5.**
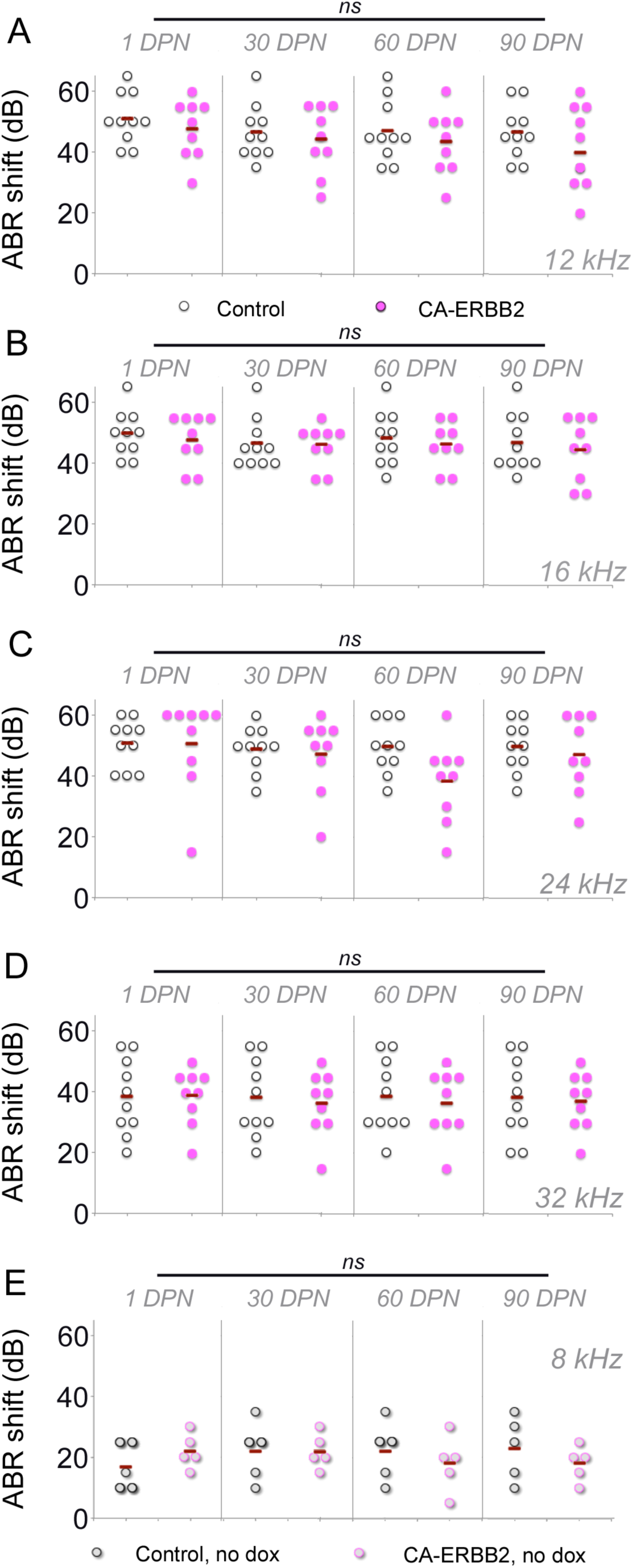
No auditory recovery is observed after CA-ERBB2 activation at higher frequencies or in the absence of dox treatment. A. ABR shifts (dB) were calculated for each mouse by subtracting original ABR thresholds from those obtained at different times after noise exposure. Positive values indicate hearing loss. ABR shifts from 10 control mice (open circles) and 9 CA-ERBB2 mice (pink circles) are plotted along with means for each group (red bars). Results for 12 kHz are shown. No recovery is observed over time for either group (p= 0.44, two-way ANOVA). B. ABR shifts (dB) for 10 control mice (open circles) and 9 CA-ERBB2 mice (pink circles) at 16 kHz. No recovery is observed for either group (two-way ANOVA p=0.83). C. ABR shifts (dB) for 10 control mice (open circles) and 9 CA-ERBB2 mice (pink circles) at 24 kHz. No recovery is observed for either group (p=0.13, two-way ANOVA). D. ABR shifts (dB) for 10 control mice (open circles) and 9 CA-ERBB2 mice (pink circles) at 32 kHz. No recovery is observed for either group (p=0.06, two-way ANOVA). E. ABR shifts (dB) for 5 control mice (open gray circles) and 5 CA-ERBB2 mice (pale pink circles) at 8 kHz, which did not receive dox treatment. This control indicates that induction of CA-ERBB2 is necessary for recovery. No recovery is observed for either group (p=0.31, two-way ANOVA).

### CA-ERBB2 transgenic mice have increased survival of hair cells

To assess hair cell loss, we dissected and immunostained whole mount cochleae and generated cochleograms from CA-ERBB2 and control littermates. Composite representative images are shown, with anti-OCM staining (Fig. 6A, B, cyan) to reveal OHCs, and anti-MYO7 (Fig. 6A, B, red) to reveal IHCs. Three regions were selected for a higher-power comparison: 45 kHz, 8 kHz, and 4 kHz (Fig. 6C, D, left, middle and right panel sets respectively). Here anti-MYO7 is in red (Fig. 6C, D, top panels), and anti-OCM in white (Fig. 6C, D, bottom panels). Figures 6C, D show that whereas, many IHCs and OHCs are missing at 45 kHz for control genotypes (Fig. 6C, left panels, red and white respectively), these cells are better preserved at 45 kHz in CA-ERBB2 mice (Fig. 6D, left panels, red and white respectively). Both genotypes have largely full complements of hair cells at 8 kHz (Fig. 6, middle panels, cf. C &D), although occasionally cells lateral to the third row of OHCs are OCM+ (Fig. 6D, middle panels). In addition, in CA-ERBB2 mice, IHCs in the apical tip express high levels of OCM (Fig. 6D, right bottom panel, cf. with C). This was observed in 3/3 CA-ERBB2 mice and in 0/4 control mice.

**Figure 6.**
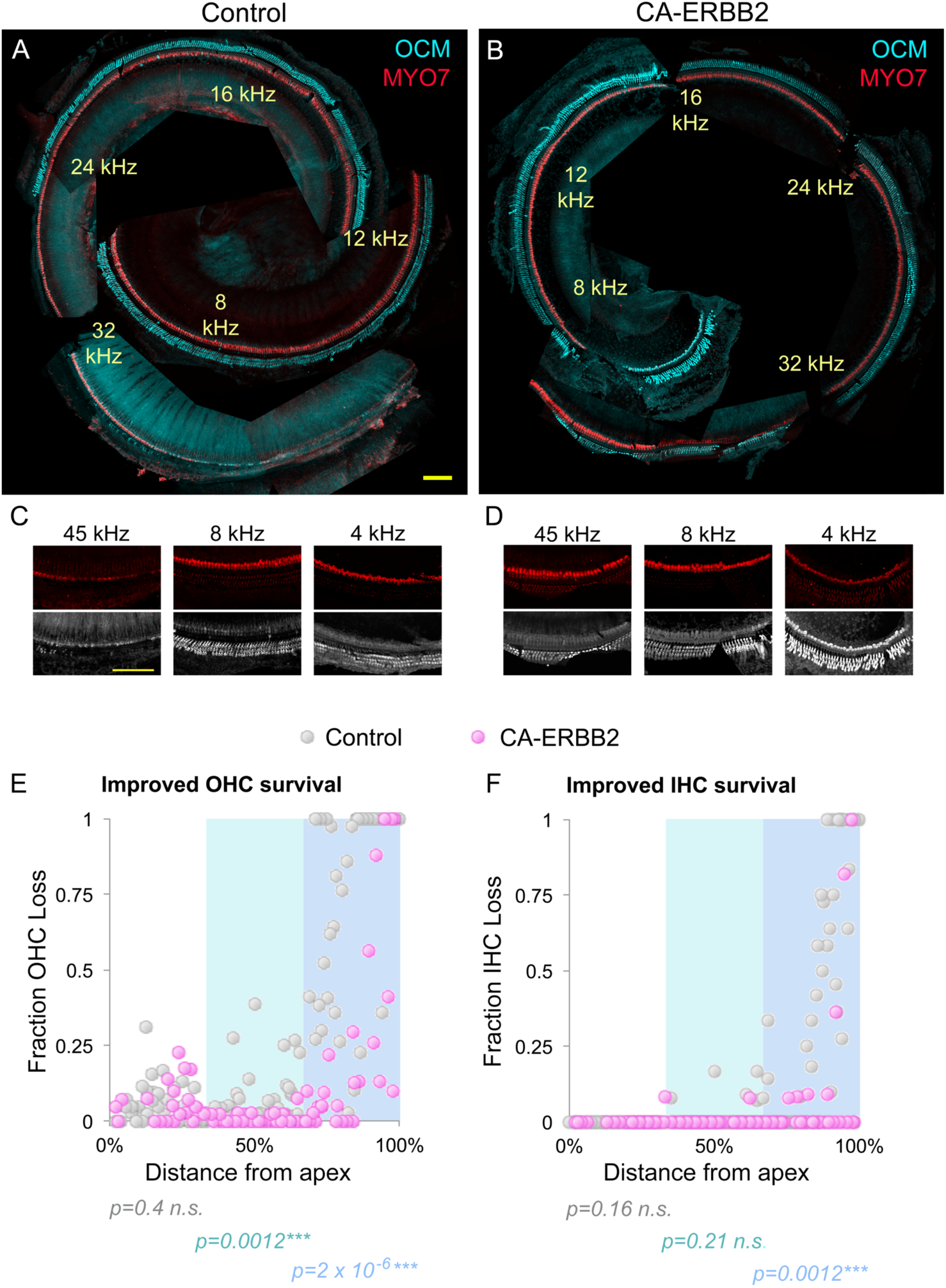
CA-ERBB2 induction correlates with improved OHC and IHC survival at higher frequencies. A. Representative control cochlea from 90 DPN, with immunostaining against OCM (cyan) to reveal OHCs and MYO7 (red) to reveal IHCs. Hair cell losses are evident in the basal segment. Frequencies are indicated in yellow. Size bar: 100 microns. B. Representative CA-ERBB2 cochlea, also from 90 DPN, same immunostaining as A. Note the HC-specific staining throughout the basal region. Frequencies are indicated in yellow. C. Anti-MYO7 (red, top) and anti-OCM (white, bottom) for 45 kHz (left), 8 kHz (middle), and 4 kHz (right) from the cochlea in (A) at higher magnification. D. Anti-MYO7 (red, top) and anti-OCM (white, bottom) for 45 kHz (left), 8 kHz (middle), and 4 kHz (right) from the cochlea in (B) at higher magnification. Note OCM+ cells lateral to the OHC (middle) and brightly OCM+ IHCs (right). E. OHC cochleogram results from control mice (gray) and CA-ERBB2 mice (pink), all at 90 DPN. Comparisons between the genotypes are made for the apical (4-16 kHz region, white), middle (16-32 kHz region, cyan) and basal (32-64 kHz region, blue) thirds of the cochlea. OHC losses were quantified in 100-micron segments along the length of each mapped organ. OHC loss is reduced in middle and basal CA-ERBB2 cochlea compared to controls (p = 0.0012 and 2 x 10^-6^ respectively, 4 control and 3 CA-ERBB2 cochleae, Kruskal-Wallis rank sum test with Bonferroni adjustment) but not in the apex (p=0.4, 4 control and 3 CA-ERBB2 cochleae, Kruskal-Wallis rank sum test with Bonferroni adjustment). F. IHC cochleogram results from control mice (gray) and CA-ERBB2 mice (pink), all at 90 DPN. Frequency regions are indicated as in (C). IHC loss is reduced in the basal region only in comparison to controls (p = 0.0012, 4 control and 3 CA-ERBB2 cochleae, Kruskal-Wallis rank sum test with Bonferroni adjustment). No differences are seen in the apical or middle regions (p = 0.16 and 0.21 respectively, 4 control and 3 CA-ERBB2 cochleae, Kruskal-Wallis rank sum test with Bonferroni adjustment).

To determine the level of hair cell survival, we quantified hair cell loss in 100-micron segments and graphed as scatter plots. Data from 3 CA-ERBB2 mice (pink circles) and 4 control mice (gray circles) were plotted together. As shown in Figure 6 E, F, basal segments in control cochleae had lost significantly more OHCs compared to CA-ERBB2 mice at 90 DPN (Fig. 6E, gray vs. pink circles, blue region, p=2 x 10^-6^, 3-4 cochleae per condition, Kruskal-Wallis rank sum test adjusted for multiple comparisons). Similarly, significantly more IHCs were lost in control cochleae than in cochlea from CA-ERBB2 mice (Fig. 6F, gray vs. pink circles, blue region, p=0.0012, 3-4 cochleae per condition, Kruskal-Wallis rank sum test adjusted for multiple comparisons). More OHC loss was also observed in control cochleae in the middle region (Fig. 6E, gray vs. pink circles, cyan region, p=0.0012, 3-4 cochleae per condition, Kruskal-Wallis rank sum test adjusted for multiple comparisons) but not in the apex (Fig. 6E white region, p=0.4, 3-4 cochleae per condition, Kruskal-Wallis rank sum test). IHC survival was not different in the middle or apical regions (Fig. 6F, p=0.21, 0.16 respectively, 3-4 cochleae per condition, Kruskal-Wallis rank sum test). OHCs in these basal regions were largely positioned in three rows above supporting cells (Fig. 6B), with some ectopic expression of OCM lateral to the OHCs. These data indicate that ERBB2 activation after damage correlates with increased OCM expression by apical cells as well as improved hair cell survival in more basal segments.

### CA-ERBB2 induces changes in morphology of synaptic ribbon in IHCs

We examined IHC morphology with immunostaining for MYO7, comparing age-matched, unexposed control mice with control mice at 90 DPN and CA-ERBB2 mice at 90 DPN (Fig. 7). 12 and 24 kHz regions were analyzed for comparison to previously published results (Kujawa and Liberman 2009, Paquette, Gilels et al. 2016, Kim, Payne et al. 2019). The morphology of IHCs were indistinguishable between the groups at 12 kHz (Fig. 7A-C, blue staining). 3D reconstructions in Amira also appeared similar (Fig. 7G-I, white). However, at 24 kHz, damaged IHCs displayed an altered appearance (Fig. 7E, cf. D). Rather than a gourd-shaped morphology filled with MYO7 (Fig. 7D), damaged IHCs in control mice displayed large gaps or holes in MYO7 staining (Fig. 7E, arrows). Gaps were also seen in CA-ERBB2 IHCs (7F, arrow). Gaps were still evident in 3D reconstructions of the data for both genotypes, where IHCs were rotated to present at the same viewing angle (Fig. 7K, L). These are similar to previous findings of long-term damage to IHCs from noise, using electron microscopy (Bullen, Anderson et al. 2019).

**Figure 7.**
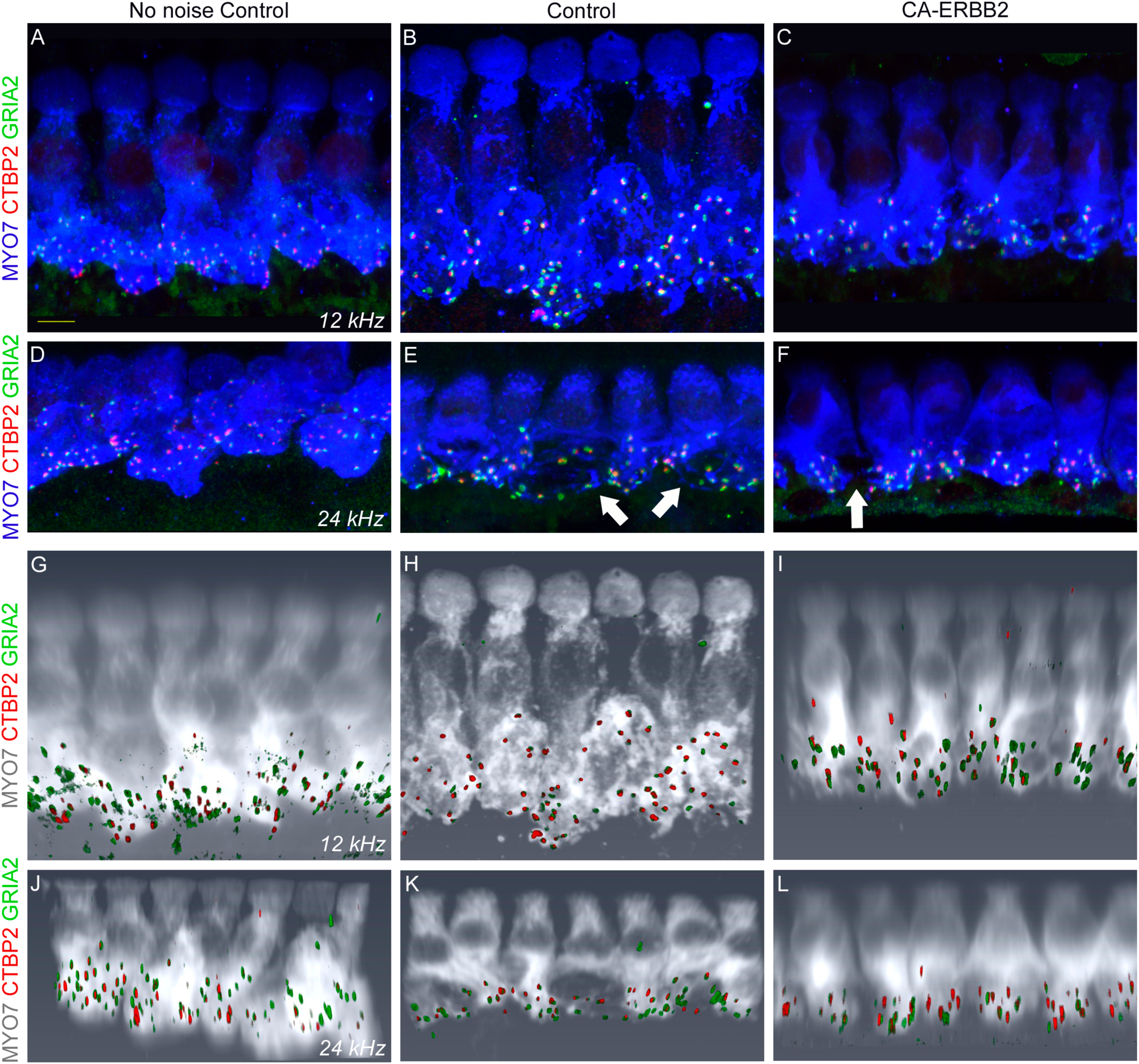
Representative IHC confocal images and Amira 3D reconstructions display synaptic labeling. A-C. Representative 200x confocal images of 12 kHz IHCs from an age-matched, control mouse without noise exposure (A), a control mouse at 90 DPN (B), and a CA-ERBB2 mouse at 90 DPN (C). Immunolabeling for CTBP2 (red), GRIA2 (green) reveal paired synaptic structures, and immunolabeling for MYO7 (blue) reveals IHC morphology. Size bar: 5 microns. D-F. Similar 200x confocal images of IHCs from the 24 kHz region, also from an age-matched, control mouse without noise exposure (D), a control mouse at 90 DPN (E), and a CA-ERBB2 mouse at 90 DPN (F). Arrows indicate IHCs with gaps in their MYO7 staining (E, F). G-I. Amira 3-dimensional reconstructions from the same stacks shown in (A-C). Strong labeling for GRIA2 is coded green, while weak labeling is in blue. CTBP2 (red) and MYO7 (white) are also shown. J-L. Amira 3-dimensional reconstructions from the same stacks shown in (D-F). IHCs were rotated to be viewed from the same approximate angle.

Noise damage can reduce synaptic pairing under conditions that impose large but temporary threshold shifts (Kujawa and Liberman 2009). Noise also promotes hypertrophy of high-frequency ribbons (Paquette, Gilels et al. 2016). We quantified synaptic pairs per IHC in Amira 3D reconstructions. The analyst was blinded to genotype, frequency, and condition. At 12 kHz, age-matched unexposed control mice, control mice at 90 DPN, and CA-ERBB2 mice at 90 DPN showed no significant differences with 12.4 ± 0.7, 14.8 ± 2.2, and 11.2 ± 1.4 synapses per IHC respectively, (mean ± s.e.m., p=0.29, ANOVA, n=3 biological replicates per condition). We plotted the ribbon sizes of 12 kHz IHCs to assess the distributions in violin plots (Fig. 8A). Mean sizes for the three conditions were 0.24 ± 0.02, 0.24 ± 0.01, and 0.38 ± 0.02 microns^3^ respectively (mean ± s.e.m., 3 biological replicates per condition). No differences were seen between control groups with or without noise and dox exposure (p=0.14, n= 3 biological replicates/260-316 synapses, Kruskal-Wallis rank sum test with multiple comparison adjustment). At 90 DPN, CA-ERBB2 IHCs had significantly larger ribbons than either control condition (Fig. 8A, p=7×10^-16^ versus age matched, unexposed control synapses, p=2×10^-14^ versus control mice at 90 DPN, 3 biological replicates/247-316 synapses, Kruskal-Wallis rank sum test with multiple comparison adjustment).

**Figure 8.**
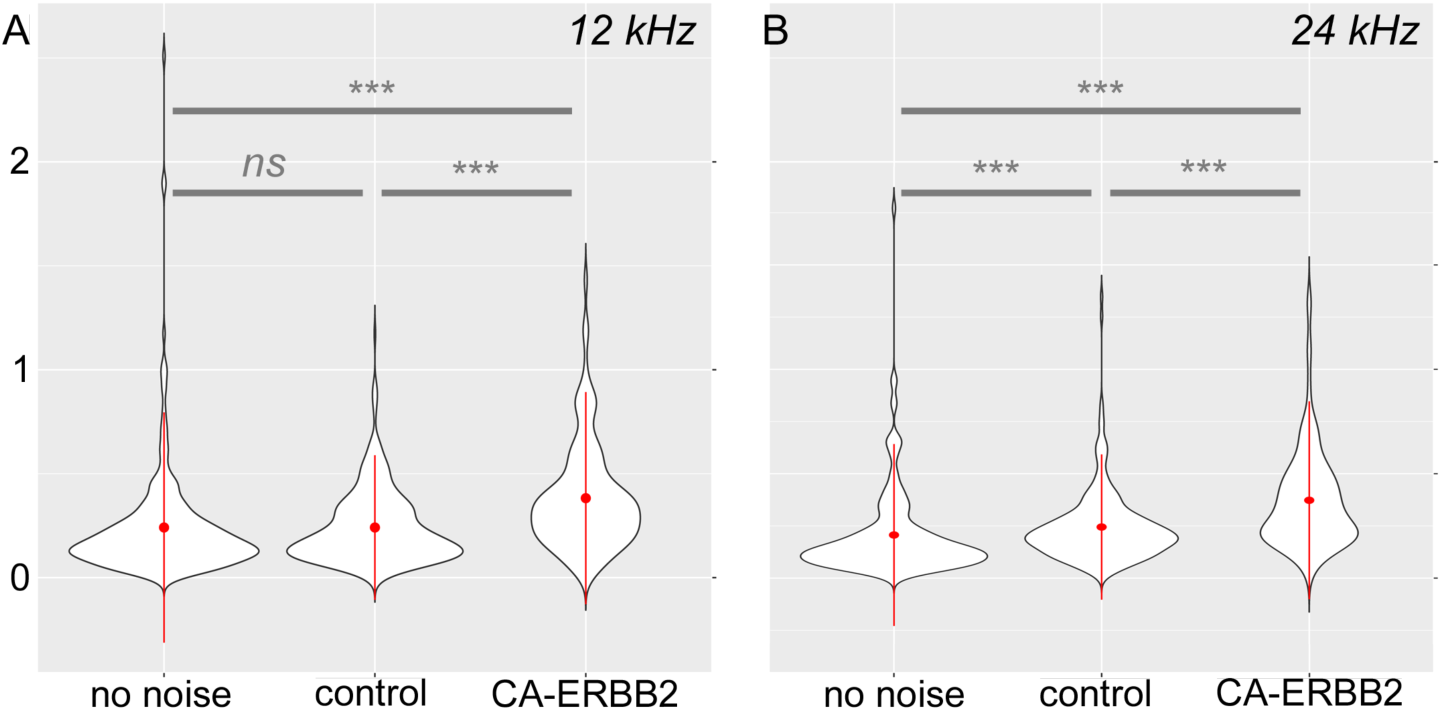
Significant differences in the synaptic ribbon size distribution are observed after CA-ERBB2 activation. A. For 12 kHz IHCs, CTBP2+ ribbons were paired with GRIA2+ post-synaptic densities in Amira 3D reconstructions as described in the text. Data from age-matched control mice without noise exposure (“no noise”) are compared to data from control mice at 90 DPN (“control”) and data from CA-ERBB2 mice at 90 DPN (“CA-ERBB2”). Approximately 240 ribbons were identified for each condition, from 6-8 IHCs per biological replicate (3 replicates each). The size distributions of paired ribbons from each condition were plotted in violin graphs. Means are represented by a red dot and standard deviations by a red bar. Shapiro-Wilk normality tests revealed the data distribution to be non-normal. The size distribution of ribbons in control mice 90 DPN is not significantly different from similar mice without noise exposure (p=0.138, Kruskal-Wallis rank sum test with multiple comparison adjustment), whereas synapses in CA-ERBB2 mice 90 DPN differ from both control mice without noise exposure and control mice at 90 DPN (p=6.6 x10^-16^, 2.1 x10^-14^, respectively, Kruskal-Wallis rank sum test with multiple comparison adjustment). B. Synaptic size distributions for 24 kHz IHCs were obtained, similarly to those at 12 kHz, and plotted in violin graphs, with means represented by a red dot and standard deviations by a red bar. Data from age-matched control mice without noise exposure (“no noise”) are compared to data from control mice at 90 DPN (“control”) and data from CA-ERBB2 mice at 90 DPN (“CA-ERBB2”). Each distribution is significantly different from each other: “no noise” vs. “control”, p=3.8 x10^-6^, “no noise” vs. “CA-ERBB2”, p=6.6 x10^-16^, and “control” vs. “CA-ERBB2”, 5.2 x10^-13^. All tests, Kruskal-Wallis rank sum test with multiple comparison adjustment, 2-3 biological replicates per condition with 6-8 IHCs per replicate.

At 24 kHz, age-matched unexposed control mice, control mice at 90 DPN, and CA-ERBB2 mice at 90 DPN also had no significant differences in the number of synapses per IHC, (12.9 ± 2.2, 10.4 ± 1.3, and 11.3 ± 1.3 synapses/IHC respectively, mean ± s.e.m., p=0.94, ANOVA, n=2-3 biological replicates per condition/164-241 synapses). Mean ribbon sizes at 24 kHz for the three conditions were 0.21 ± 0.02, 0.24 ± 0.01, and 0.37 ± 0.02 microns^3^ respectively (mean ± s.e.m., 2-3 biological replicates per condition). Noise exposed control IHCs had larger ribbons than age-matched unexposed control IHCs (Fig. 8B, p=4×10^-6^, n= 2-3 biological replicates/164-241 synapses, Kruskal-Wallis rank sum test with multiple comparison adjustment). Noise exposed CA-ERBB2 IHCs were significantly larger than both control conditions (Fig. 8B, p=7×10^-16^ versus age matched, unexposed control synapses, p=5×10^-13^ versus control mice at 90 DPN, 2-3 biological replicates/164-241 synapses, Kruskal-Wallis rank sum test with multiple comparison adjustment). Taken together, we show no significant synaptic loss in this experimental system, but do suggest that after CA-ERBB2 induction, some intracellular processes within IHCs are altered in a way that leads to larger synaptic ribbon sizes. It is noted that these comparisons were made outside the region that exhibited functional recovery.

We sought to better characterize the “gaps” in anti-MYO7 staining observed in IHCs (Fig. 7) after damage, using cryosections (Fig. 9). Since MYO7 is an actin-binding protein, we compared anti-MYO7 staining patterns (Fig. 9, green) to anti-MAP2, a microtubule-associated protein in the organ of Corti (Oshima, Okabe et al. 1992), Fig. 9, red). To outline the IHCs, we included immunostaining against GLAST/EAAT1, a glutamate transporter expressed in interphalangeal supporting cells (Chen, Kujawa et al. 2010), Fig. 9, magenta). We found that areas within the IHCs where anti-MYO7 staining was reduced (Fig. 9, arrows) also had reduced MAP2 immunostaining (Fig. 9, arrows), although precise co-expression was infrequent. This suggests that gaps in anti-MYO7 staining could reflect local injury to IHC cytoskeleton. Cytoskeletal gaps were irregular in shape, and typically near the synapses, although some were observed in perinuclear regions (Fig. 9A) or between the nucleus and stereocilia (Fig. 9C). There was no obvious difference between the genotypes, suggesting that the persistence of cytoskeletal gaps is not affected by ERBB2 activation.

**Figure 9.**
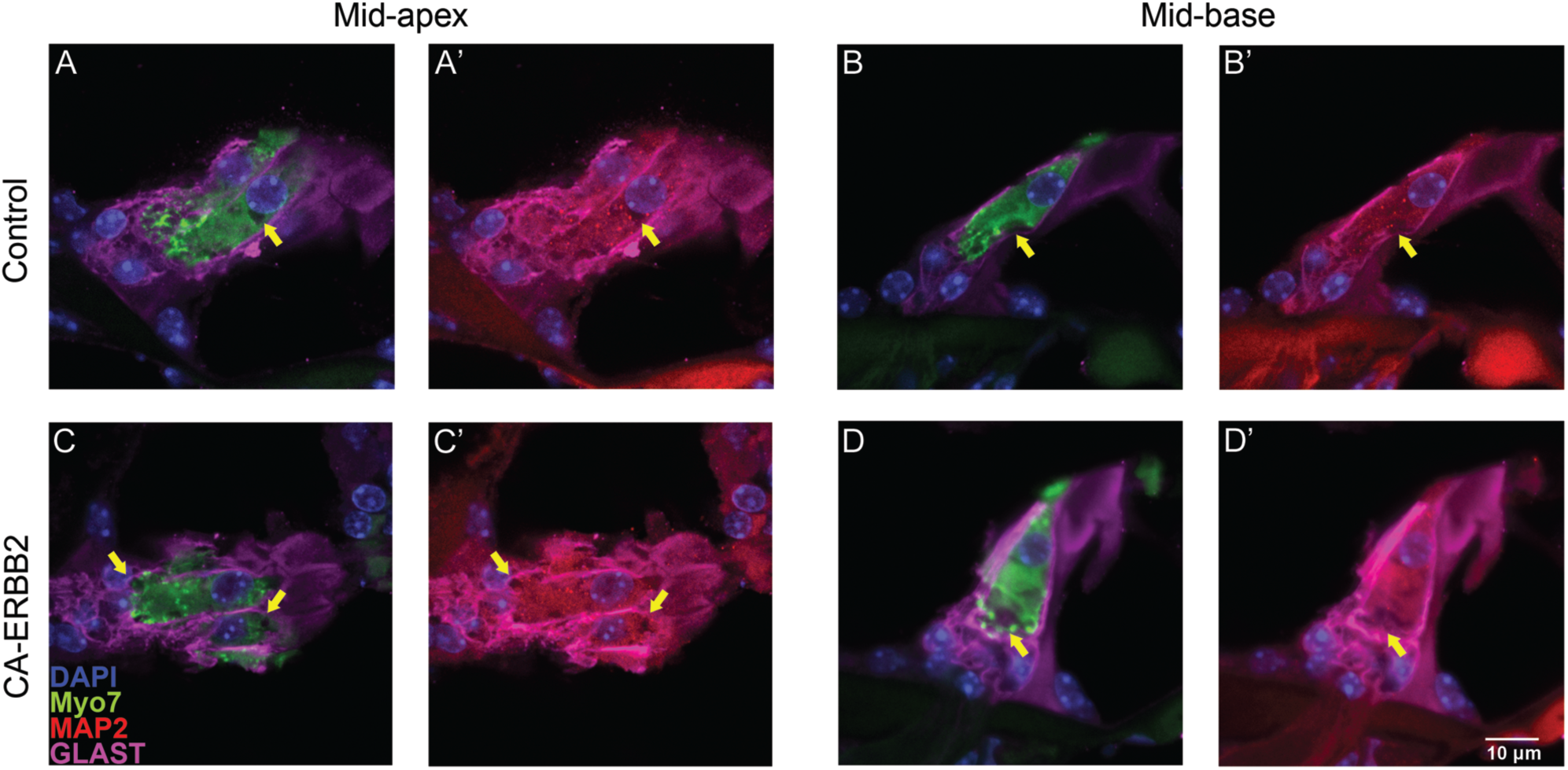
Loss of MYO7 immunoreactivity in IHC bodies may reflect damage to the cytoskeleton. A, A’. A single optical section through the center of a cryosection of a IHC immunostained against MYO7 (A, green), MAP2 (A’, red), GLAST (A, A’, magenta) and co-stained with DAPI (A, A’, blue). The IHC is from the mid-apical region of a control mouse cochlea obtained at 90 DPN. A staining “gap” is highlighted next to the nucleus (arrow). B, B’. An optical section from the center of a similarly stained IHC, from the mid-basal region of a control mouse cochlea obtained at 90 DPN. A large gap is highlighted (arrow). C, C’. An optical section from the center of a similarly stained IHC, from the mid-apical region of a 90 DPN cochlea in which CA-ERBB2 had been activated. Gaps in MYO7 (C, green) and MAP2 (C’, red) are also evident. D, D’. An optical section from the center of a similarly stained IHC, from the mid-basal region of a 90 DPN cochlea in which CA-ERBB2 had been activated. A large gap in cytoplasmic staining is highlighted (arrow). Images were chosen to illustrate gaps in specific places in IHCs from 3 biological replicates of each genotype. Size bar: 10 microns.

### CA-ERBB2 induction is not sufficient to significantly improve morphology of stereocilia in hair cells

We examined stereociliary morphology in CA-ERBB2 and control cochleae at 90 DPN (Fig. 10). In mice without noise or dox exposure, IHC stereocilia at 12 kHz have tapered bases near the cuticular plate (Fig. 10A). Blebs, as evidenced in this image, are fixation artifacts (Fig. 10A). OHC stereocilia at 12 kHz in the same cochleae are precisely organized, reflecting OHC organization (Fig. 10B). In damaged, control cochleae at 8-12 kHz, IHC stereocilia have normal morphology but appear sparse (Fig. 10C). At more basal regions (32 kHz) some IHC stereocilia are fused together (Fig. 10E, yellow arrow). OHC stereocilia were comparable in morphology to non-damaged, age-matched controls at 8-12 kHz (Fig. 10D), and missing OHCs were observed in scans at 40 kHz (Fig. 10F). Three months after CA-ERBB2 induction, fused IHC stereocilia were also evident at 8 kHz (Fig. 10G, right arrow) and some stereocilia were mis-oriented (Fig. 10G, left arrow, 10I, arrow). However, compared to control apical IHCs, CA-ERBB2 appeared to have a more complete complement of stereocilia (Fig. 10G, cf. with C). At 48 kHz, IHCs with missing and stunted stereocilia were observed (Fig. 10I, brackets, note comparable 2-micron size bar to the same in 10G). No ectopic OHC stereocilia were observed at 8-12 kHz in CA-ERBB2 cochleae, and stereocilia morphology was similar to control noise damaged cochleae (Fig. 10H, cf. 10D). In CA-ERBB2 mice, 60 kHz OHCs also exhibited stunted and fused stereocilia (Fig. 10J, arrows). We quantified IHC bearing fused stereocilia in undamaged, control damaged, and CA-ERBB2 damaged cochleae, finding 0/98 (0%), 4/41 (9.8%), and 20/93 (21.5%) respectively. These data suggest that severe damage to the cochlea persistently alters stereociliary morphology, but that these processes are not improved by CA-ERBB2 induction. However, loss of IHC stereocilia likely impacts cochlear function, and control noise-damaged apical IHCs appeared to have fewer stereocilia compared to noise damaged IHCs after CA-ERBB2 induction.

**Figure 10.**
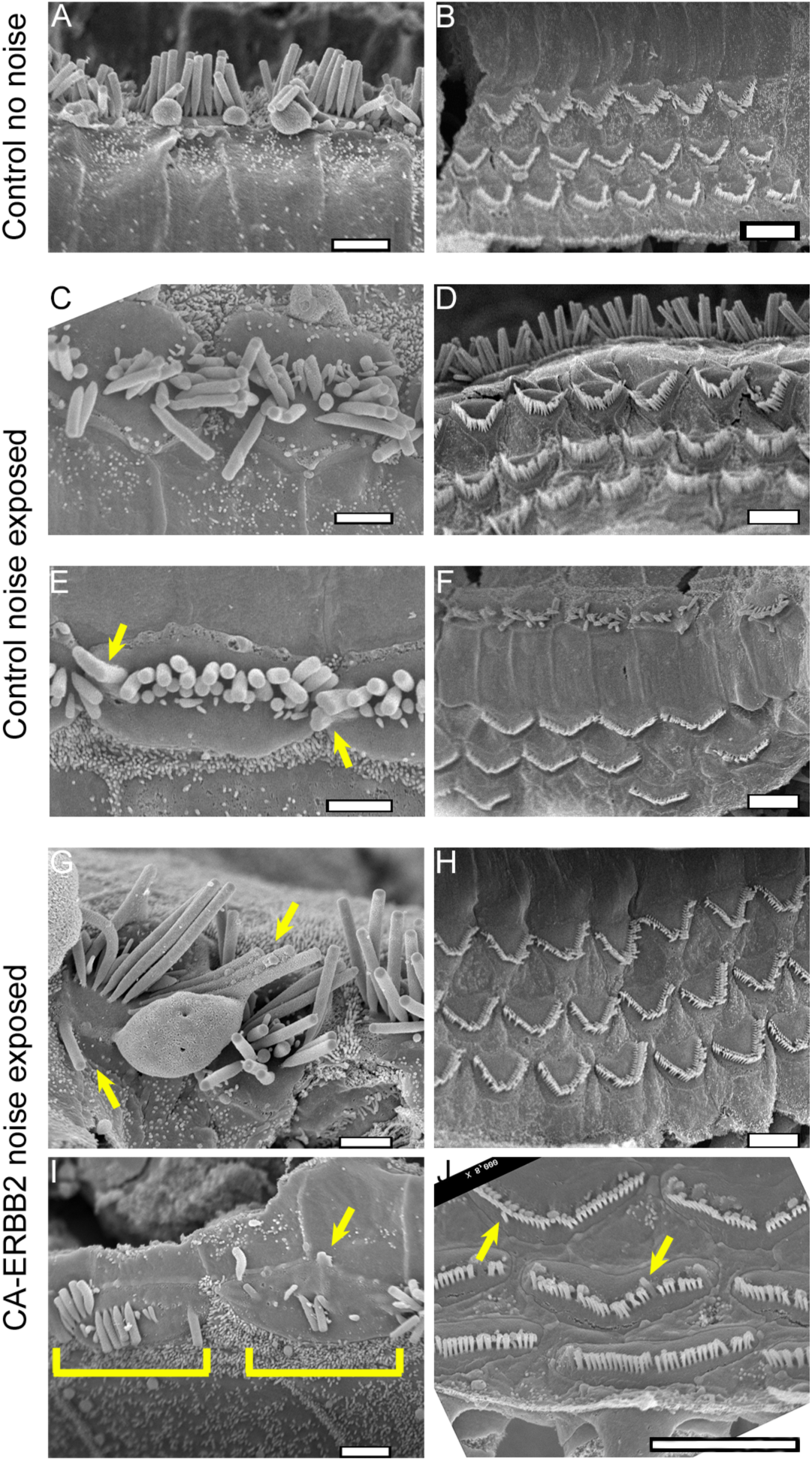
Persistent stereociliary defects are seen in both control and CA-ERBB2 cochleae. A, B. Stereocilia in in an age-matched control cochlea with no history of noise or dox exposure. 12 kHz IHC stereocilia (A) show a characteristic tapered morphology near the cuticular plate. Rounded blebs are a fixation artifact. 12 kHz OHC stereocilia (B) reflect regular OHC patterning. C-F. IHC (C, E) and OHC (D, F) stereocilia in a control mouse euthanized at 90 DPN. 8-12 kHz IHCs are missing stereocilia (C). At 32 kHz, IHCs have fused stereocilia (E arrow). OHC stereocilia appeared grossly normal at 8-12 kHz (D). Missing OHCs were evident at 32 kHz (F). G-J. IHC (G, I) and OHC (H, J) stereocilia in a CA-ERBB2 mouse euthanized at 90 DPN. Mis-oriented (G, left arrow) and fused (G, right arrow) stereocilia were seen at 8 kHz in CA-ERBB2 cochleae; however, their numbers appear largely intact. At 45 kHz, IHCs with stunted and missing stereocilia (I, brackets) were observed. OHC stereocilia at 8-12 kHz in CA-ERBB2 cochleae (H) were comparable to no damage controls (cf. B) and noise-damaged controls (cf. D). Fused stereocilia (J, arrows) were observed in CA-ERBB2 OHCs at 60 kHz. Left column size bars: 2 microns. Right column size bars: 5 microns.

## Discussion

NIHL is a lifelong disability partially characterized by raised auditory thresholds and OHC loss. Here we tested the hypothesis that activating ERBB2 signaling in supporting cells after noise and doxycycline exposure can mitigate cochlear damage. Using a genetic expression system to drive a constitutively active ERBB2 transgene, we show that this correlates with a delayed, partial recovery of hearing function in the lower frequency of young adult mice after traumatic noise exposure. While recovery was only seen at 8 kHz, it was comparable in extent to another report using post-damage treatments (Du, Cai et al. 2018) and better than earlier attempts (Mizutari, Fujioka et al. 2013). Basal OHCs and IHCs, as well as middle OHCs have significantly greater survival in mice with CA-ERBB2 activation. Apical IHCs express higher levels of OCM after CA-ERBB2 expression, and may retain more stereocilia compared to control IHCs. Mid-frequency auditory synapses are maintained in this noise-damage paradigm. Synaptic ribbons in both 12 and 24 kHz regions are significantly hypertrophied in cochleae with induced CA-ERBB2 expression. IHC cytoskeletal and stereociliary damage both persist in CA-ERBB2 expressing and control cochleae. No ectopic HCs were observed by SEM. Taken together, we propose that activation of this pathway in cochlear supporting cells may partially restore hearing after noise damage in mammals through an unknown mechanism. The restriction of putative CA-ERBB2-expressing cells to the apical region even though damage was mitigated throughout the cochlea suggests that part of this mechanism is systemic or diffusible.

We believe that this is the first report showing reversal of permanent auditory threshold shifts in mammals over a time course similar to auditory recovery from noise in avians. We observed partial but significant improvements in hearing thresholds in noise damaged mice after CA-ERBB2 induction (Fig. 3). This recovery was only at the lowest frequency tested, 8 kHz (Fig. 3), and required doxycycline administration (Fig. 5). After CA-ERBB2 induction, wave I latencies at 8 kHz were slightly but significantly delayed compared to control, noise damaged littermates (Fig. 4). As CA-ERBB2 expression was likely confined to the apical third of the cochlea (Fig. 1), hearing restoration may require proximity to cells that induce ERBB2 activity.

We also found that there are significantly more preserved OHCs and IHCs in basal cochleae of noise-damaged mice after CA-ERBB2 induction (Fig. 6). Basal OHCs and IHCs are not disorganized or ectopic (Fig. 6), supporting cell loss was not observed, and no ectopic stereocilia were observed (Fig. 10). Thus, we interpret these results to mean that CA-ERBB2 signaling in supporting cells likely promotes recovery and survival of damaged hair cells rather than the regeneration of new hair cells. In the mouse utricle, supporting cells respond to damaged hair cells by secreting HSP70, promoting their survival (May, Kramarenko et al. 2013). A similar mechanism may be at work here. It is remarkable that this effect was seen at a significant distance from the regions where CA-ERBB2 expression is predicted to have occurred. We note that noise damages the cochlea in a basal to apical gradient, with the greatest cellular losses in the high-frequency, basal regions. Auditory function is lost throughout most of the noise-damaged cochlea, and any residual function confined to the low-frequency, apical regions. It is promising that both low-frequency auditory function and higher-frequency hair cell survival were improved in cochleae after CA-ERBB2 expression, consistent with the interpretation that protection occurred throughout the organ. Our data suggest that damaged and dysfunctional hair cells might be repaired to restore function in the apical cochlea.

We report on the number of synapses observed on IHCs at 12 and 24 kHz. The synapse range as defined by (Kujawa and Liberman 2009) at the higher end with ∼18 synapses per IHC, and by (Cederroth, Park et al. 2019) at the lower end with 13-15 synapses per IHC, places our results within the lower end. We do not find differences in the numbers of synapses on IHCs when comparing CA-ERBB2 mice exposed to noise, control littermates exposed to noise, or control littermates without noise exposure. Pure CBA/CaJ mice permanently lose synapses after a noise exposure that produces a temporary hearing loss (Kujawa and Liberman 2009). Similar results were reported for FVB/nJ mice (Paquette, Gilels et al. 2016) and rhesus monkeys (Valero, Burton et al. 2017). In contrast, both guinea pigs (Liu, Wang et al. 2012) and C57BL6/J mice (Kim, Payne et al. 2019) have been reported to repair some damaged synapses. The latter report may be relevant to our studies, as our mice are F1 hybrids of C57BL6/J and CBA/CaJ.

We report significant ribbon hypertrophy at both low- and mid-range frequencies after noise damage and CA-ERBB2 activation (Fig. 8). The analysis was performed outside the region that shows functional recovery, but allows us to make comparisons with previously published results. Sustained hypertrophy of mid-frequency, e.g. 24 kHz, ribbons is observed in control mice after noise exposure (Fig. 8B), both with and without permanent threshold shift (Paquette, Gilels et al. 2016) and in multiple mouse lines (Kim, Payne et al. 2019). Hypertrophy occurs in the absence of synaptic loss, meaning that noise damage causes ribbons to grow. CTBP2 proteins exchange dynamically, but slowly, in hair cells, with a half-life in ribbon structures of around 2 hours (Chen, Chou et al. 2018). In zebra fish, the enlargement of ribbons reduces spontaneous spike rates and alters stimulus encoding latencies (Sheets, He et al. 2017). Thus, a mechanism that maintains ribbon size likely facilitates coordinate signaling and possibly sound encoding (Carney 2018). In zebrafish, reducing Ca(V)1.3 activity enlarges hair cell ribbons, and the enlargement of ribbons leads to larger calcium currents (Sheets, Kindt et al. 2012). These suggest that calcium currents can regulate ribbon size. To get a long-lasting enlargement of ribbons, noise damage could suppress calcium currents, or make the ribbon less sensitive to calcium by altering a putative internal IHC feedback mechanism. The second mechanism could involve modifications in levels or activities of calcium binding proteins associated with the ribbon, such as ArfGAPs (Dembla, Wahl et al. 2014) or otoferlin. It will be interesting to see if ribbon size could also impact wave I latency (Fig. 4), or if higher levels of OCM alter ribbon sizes (Fig. 6). We do not assert that ribbon hypertrophy drives hearing recovery. Rather, it may reflect intrinsic changes in the IHC after noise damage; this could affect its responsiveness to stimuli. Further studies are needed to better understand synaptic maintenance, structure, and responsiveness to damage, especially at 8 kHz where functional recovery is evident.

One major outstanding question in otolaryngology is the relative noise contributions of different noise damage correlates to hearing loss. Cochlear function requires multiple cell types to work together, including OHCs, IHCs, supporting cells, spiral ganglion neurons, efferent projections, and potassium recycling cells. We have focused attention on inner hair cells, their synapses, their cytoskeleton, and their stereocilia (Fig. 7-10), which show clear and persistent signs of damage. We observed two differences in apical IHCs after CA-ERBB2 induction that might correlate with improved function. First, there was a wider distribution of OCM expression in the apical cochlea after CA-ERBB2 induction, including OCM+ IHCs (Fig. 6). A recent report using single cell sequencing has shown that apical IHCs do express low levels of OCM (Tang, Chen et al. 2019), suggesting that this represents an up-regulation. Changes in Ca++ regulation are a potential mechanism for repair. Secondly, while 8 kHz IHC stereocilia exhibited significant abnormalities in morphology at 8 kHz, more stereocilia appeared to be present per cell (Fig. 10). Stereocilia loss is a plausible explanation for loss of function. Notably, we saw no differences in synaptic sizes between noise damaged controls and no noise controls at 12 kHz, even though very little hearing was observed at 12 kHz, suggesting that this measurement is less relevant to threshold shifts. Further research is needed to understand which aspects of noise damage are critical for function and which ones are correlative.

Future experiments will address several issues that we were unable to resolve for this publication. Ongoing experiments seek to identify mechanisms of hearing recovery, which may be due to changes to efferent projections and/or supporting cell functions in addition to repair of damaged hair cells. These must also be explored at 8 kHz, where functional recovery is observed. Second, ERBB2 phosphorylation may be modulated differently in adult supporting cells compared to neonatal supporting cells, as we were unable to directly confirm expression of CA-ERBB2 through the detection of p-ERBB2 protein as we did in a previous report (Zhang, Wang et al. 2018) Understanding the stability of p-ERBB2 is a critical feature for ongoing experiments. Third, it will be interesting to determine if synaptic changes downstream of CA-ERBB2 activation require noise damage. Lastly, experiments to assess both the proliferation of supporting cells after CA-ERBB2 induction and to identify ERBB2’s downstream mediators are ongoing.

We acknowledge some limitations in our methods. First, to assess the location of putative CA-ERBB2+ cells, we tracked the expression of GFP from the internal ribosomal entry site, as pERBB2 immunoreactivity was not observed. We indirectly demonstrate ERBB2 activation by detecting transcripts for the transgene (S. Fig. 1) and performing no-dox controls, where no improvement was seen (Fig. 5). Second, the variance in CA-ERBB2 recovery at 8 kHz was great enough that we could not show statistically significant improvement when only CA-ERBB2 mice were considered; the improvement described is between the two genotypes. This may be due to variability in the distribution or number of CRE-expressing cells. Lastly, genetic models such as the ones we use here may harbor strain variants linked to genetic modifications that could affect results. While CA-ERBB2 mice and control littermates had no differences in hearing thresholds prior to noise damage (Fig. 2), we did find that CA-ERBB2 mice had significantly higher wave I amplitudes (Fig. 4), suggesting a potential genetic difference. The commonly used 129SvEv strain used to generate most transgenic mice has a well-known resistance to noise damage (Yoshida, Hequembourg et al. 2000). Those variants affect permanent threshold shifts, not the delayed recovery as in assays presented here. We included controls, such as the no dox control, to support our interpretation that ERBB2 activation is the source of the recovery. Nonetheless, as more studies are conducted on mice months after large permanent threshold shifts, it is possible that strain differences may emerge in the propensity for hearing restoration.

Previous reports have shown that CA-ERBB2 can activate the downstream effector PI3K (Xie, Chow et al. 1999, Zhang, Wang et al. 2018). While it is attractive to imagine that endogenous ERBB2 activation could confer detection of stretch damage (Vermeer, Einwalter et al. 2003) in cochlear epithelium, an alternative explanation holds that transgenic CA-ERBB2 over-expression artificially activates PI3K to drive hearing restoration. Previous studies have implicated PI3K in cochlear protection from ototoxicity. Pharmacological blockade of PI3K increases OHC death from gentamicin in the neonatal rat organ of Corti (Chung, Pak et al. 2006), and genetic activation of PI3K through the deletion of its inhibitor, PTEN, reduces hair cell death from cisplatin (Jadali, Ying et al. 2017) or gentamicin toxicity (Jadali and Kwan 2016). Under this interpretation, our work extends these findings in several important ways. First, we show that activation in supporting cells alone is sufficient to promote hair cell survival, as the previous studies used either pharmacological inhibitors, deleted PTEN in both supporting cells and hair cells, or both (Jadali and Kwan 2016, Jadali, Ying et al. 2017). Second, our work indicates that the induction of CA-ERBB2, and/or its potential effector PI3K, can rescue adult hair cells from a damage-induced death. Third, we note that CA-ERBB2 is induced three days after noise damage, indicating a substantial window of time for therapeutic intervention after noise exposure. Lastly, the time course of hearing restoration suggests this signal initiates a slow repair process that partly reverses permanent threshold shifts. Therapies incorporating technology based on this finding could be bundled with other methods, such as those that reduce immediate damage, for combinatorial effects.

## Conclusions

Young adult mice with noise induced hearing loss partially recover their low-frequency hearing after ERBB2 signaling is artificially activated in a small number of cochlear apical supporting cells. This recovery manifests 60 days after damage, and is maintained for at least another 30 days. It correlates with significantly increased retention of high-frequency IHCs and OHCs, suggesting mitigation of a gradient of damage along the length of the cochlea. Assessments of IHC differences between genotypes reveal a potentially greater complement of stereocilia in CA-ERBB2 8 kHz IHCs and the up-regulation of a calcium regulator, OCM. Downstream intermediaries of ERBB2 signaling are unknown, but possibly diffusible in nature. This is the first report to assess reversals of permanent hearing loss from noise in mice.

## Declaration of interest statement

Drs. Zhang and White are inventors on a patent, “ERBB3 activators in hearing restoration” which is related to this work.

## Author Contributions

J. Zhang and P. M. White conceived the experimental design; J. Zhang, D. Na, H. J. Beaulac, M. Dilts, A. Bullen, and P. M. White performed experiments; J. Zhang, A. Bullen, K. S. Henry and P. M. White performed analysis; and P. M. White wrote the first draft of the paper. All authors contributed to subsequent edits and approved the final version.

## Acknowledgements

We gratefully acknowledge Dr. Anne Luebke, who maintains the URMC Small Animal Auditory Testing Core; Dr. John Ashton from the URMC Genomics Core for the qPCR studies, Dr. V. Kaye Thomas, who maintains the URMC Center for Advanced Light Microscopy and Nanoscopy; Dr. Jian Zuo for the *Fgfr3-*iCre mouse strain; Dr. Lin Gan for the ROSA-floxed rtTA/GFP mouse strain; Dr. Dorota Piekna-Przybylska for critical reading, and Ms. Lisa Shah, for technical assistance.

## Data Accessibility Statement

Upon acceptance of this manuscript, all original images, ABR waveforms, and analysis will be deposited at the UR Research Institutional Repository at the University of Rochester.

---

**Supplemental Figure 1.**
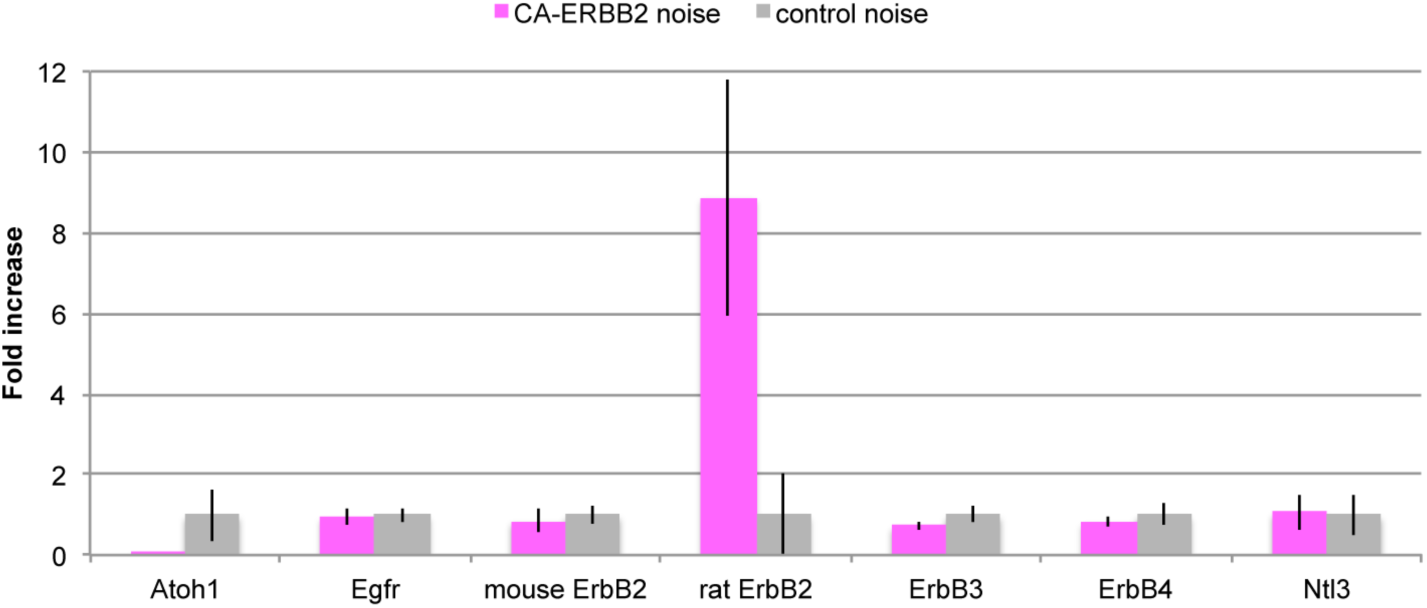
Quantitative PCR for selected transcripts from cochleae at 7 DPN. CA-ERBB2 (pink) and control (gray) littermates were exposed to noise, later receiving injections for doxycycline and furosemide at 3 DPN. At 7 DPN, the mice were euthanized and mRNA was extracted from their cochleae. Quantitative PCR was performed for seven mouse genes (Atoh1, Egfr, ErbB2, ErbB3, ErbB4, and Ntl3) as well as for rat ErbB2, which detects the CA-ERBB2 transgene. Results were normalized to the geometric mean of Gapdh and Hprt1 in the same run. Only the transgene was significantly different between the genotypes (p=0.027 for rat ErbB2, two-way Student’s t-test with Bonferroni adjustment, n=4-5 cochlea).

## Notes

**Support:** This work was funded by the National Institute of Health R01 DC014261, and by grants from the Schmitt Foundation and UR Ventures.

### Competing Interest Statement

The authors have declared no competing interest.

